# Pan-cancer organoid validation of tumor outlier chromosomal amplification events

**DOI:** 10.1101/2021.10.05.463147

**Authors:** Ameen A. Salahudeen, Kanako Yuki, Jose A. Seoane, Amanda T. Mah, Amber R. Smith, Kevin Kolahi, Sean M. De la O, Daniel J. Hart, Jie Ding, Zhicheng Ma, Sammy Barkal, Navika D. Shukla, Chuck Zhang, Michael A. Cantrell, Arpit Batish, Tatsuya Usui, David Root, William Hahn, Christina Curtis, Calvin J Kuo

## Abstract

Somatic copy number gains are pervasive in many cancer types, yet their roles in oncogenesis are often poorly explored. This lack of understanding is in part due to broad extensions of copy gains across cancer genomes spanning large chromosomal regions, obscuring causal driver loci. Here we employed a multi-tissue pan-organoid modeling approach to validate candidate oncogenic loci identified within pan-cancer TCGA data by the overlap of extreme copy number amplifications with extreme expression dysregulation for each gene. The candidate outlier loci nominated by this integrative computational analysis were functionally validated by infecting cancer type-specific barcoded full length cDNA lentiviral libraries into cognate minimally transformed human and mouse organoids bearing initial oncogenic mutations from esophagus, oral cavity, colon, stomach, pancreas and lung. Presumptive amplification oncogenes were identified by barcode enrichment as a proxy for increased proliferation. Iterative analysis validated *DYRK2* at 12q15, encoding a serine-threonine kinase, as an amplified head and neck squamous carcinoma oncogene in *p53*^*-/-*^ oral mucosal organoids. Similarly, *FGF3*, amplified at 11q13 in 41% of esophageal squamous carcinomas, was validated in *p53*^*-/-*^ esophageal organoids in vitro and in vivo with pharmacologic inhibition by small molecule and soluble receptor FGFR antagonists. Our studies establish the feasibility of pan-organoid contextual modeling of pan-cancer candidate genomic drivers, enabling oncogene discovery and preclinical therapeutic modeling.

## INTRODUCTION

Somatic copy number aberrations (SCNAs) in the form of amplifications or deletions are a common genomic event in solid tumors (Beroukhim et al., 2010; Zack et al., 2013) and have been successfully targeted by therapeutics such as trastuzumab (Oh and Bang, 2020), whereas deletion events, such as *MTAP*, are currently the subject of therapeutic development for synthetic lethal interactions (Kryukov et al., 2016). However, a comprehensive understanding of a given SCNA’s contribution to tumor biology and patient outcomes remains an aspirational goal (Kristensen et al., 2014). Furthermore, while recurrent amplified or deleted regions are increasingly delineated with large scale genomic studies, many being prognostic, which genes within the amplicon contribute to oncogenesis is often unknown.

Several methods have been developed for the identification of recurrent somatic copy number alterations (SCNAs), including GISTIC2, which enables the identification of focal alterations (Beroukhim et al., 2010; Zack et al., 2013) and RUBIC (van Dyk et al., 2016). However, copy number amplicons are often broad, spanning multiple megabases, making it challenging to pinpoint the driver. In order to identify loci within SCNA, we identify candidate “outlier” genes demonstrating both high-level copy number alterations and concordant extreme expression dysregulation. The outlier approach is highly discriminatory, prioritizing candidate drivers within broad SCNA regions, highlighting known oncogenes and tumor suppressors and numerous novel candidates, as demonstrated in breast cancer (Curtis et al. 2012), several of which have been functionally validated (Sanchez-Garcia et al., 2014; Turner et al., 2010). Related methods have been used to identify chromosomal rearrangements (Tomlins et al., 2005) by comparing “normal” with dysregulated gene expression. However, to date a systematic screen of candidate SCNA drivers across primary tissues not been undertaken.

To functionally test hypotheses from genomic cancer analyses, immortalized cell lines or xenograft models are frequently used. Such systems are not optimal for several reasons including mono- or oligo-clonality (i.e. lack of tumor heterogeneity), secondary mutation burden in cell lines, the limited tractability of patient derived xenografts, and low throughput of genetically engineered mice. In contrast, in vitro organotypic culture of untransformed primary tissues as 3-dimensional organoids offers an extremely promising approach for driver oncogene validation (Lo et al., 2020). Indeed, organoids faithfully recapitulate multilineage differentiation and tissue architecture, and yet retain experimental tractability for in vitro genetic and pharmacologic studies (Broutier et al., 2017; Du et al., 2020; Francies et al., 2019; Huang et al., 2015; Li et al., 2018; Li et al., 2014; Matano et al., 2015; Nadauld et al., 2014; Salahudeen and Kuo, 2015; Sato et al., 2011; van de Wetering et al., 2015). We and others have initiated gastrointestinal malignancies by oncogene-engineering wild-type organoids from mouse (Li et al., 2014; Nadauld et al., 2014) and human tissues (Drost et al., 2015; Lo et al., 2021; Matano et al., 2015). Importantly, these cancer organoid models are generated by introduction of “first hit” oncogenic alleles into a normal wild-type genome, representing predominant drivers of the cognate cancer type. Subsequently, putative oncogenic loci can be overlaid and functionally assessed in a scalable, rapid, and reproducible manner.

Here, we pursued a pan-cancer approach to amplified oncogene validation, exploiting the *tabula rasa* background of a diverse range of first hit-engineered organoid models to rapidly interrogate the oncogenic potential of candidate amplified/overexpressed outlier loci across several solid tumor histologies. The transduction of lentiviral barcoded open reading frame (ORF) libraries, representing tumor subtype-specific SCNAs, into cognate tissue specific organoid models, then allowed systematic functional screening of oncogenicity, followed by iterative hit validation and assessment of therapeutics.

## RESULTS

### Pan-cancer bioinformatic identification of tumor outlier loci exhibiting matched extreme copy number alteration and expression

The analysis of copy number (CN) somatic events in cancer is impeded by broad extension of somatic CN events over the genome, thus increasing the likelihood of false positives. Most methods for copy number driver identification analyze only DNA and thus include alterations that have no effect on expression. The approaches that use an integrative expression/copy number methodology often include the expression information by identifying genes whose expression is correlated with CN values. This approach has the limitation that regression between the CN and expression values is often polynomial, and not evenly distributed between high and low copy number values (i.e. coefficient between higher copy number and higher expression is always higher than the lower copy number and lower expression), thus potentially including false negatives. Building upon our prior studies (Curtis et al., 2012), we sought to integrate gene expression information into our model to more accurately identify outliers where we propose putative drivers are those with a modification in the copy number landscape at given position with a corresponding functional effect in gene expression.

To refine putative amplified SCNAs consequential to oncogenesis, we matched gene-based extreme copy number events (amplifications and deletions) with extreme expression effects (**Fig. 1A**). In order to identify which samples falls in the overexpressed tail, we calculate the threshold for the 5% right and 5% left values in the theoretical distribution (represented by the vertical lines **Fig. S1A**). With these two thresholds we identified expression outliers, which are the values in the real expression distribution that are greater than right threshold (overexpression outliers). We then matched overexpression outliers with extreme copy number amplifications (defined as samples with a segmented copy number value higher than 6 times the standard deviation) in sequencing data from TCGA datasets COAD, ESCA, HNSC, LUAD, PAAD and STAD (**Fig. 1B-G**).

**Figure 1.**
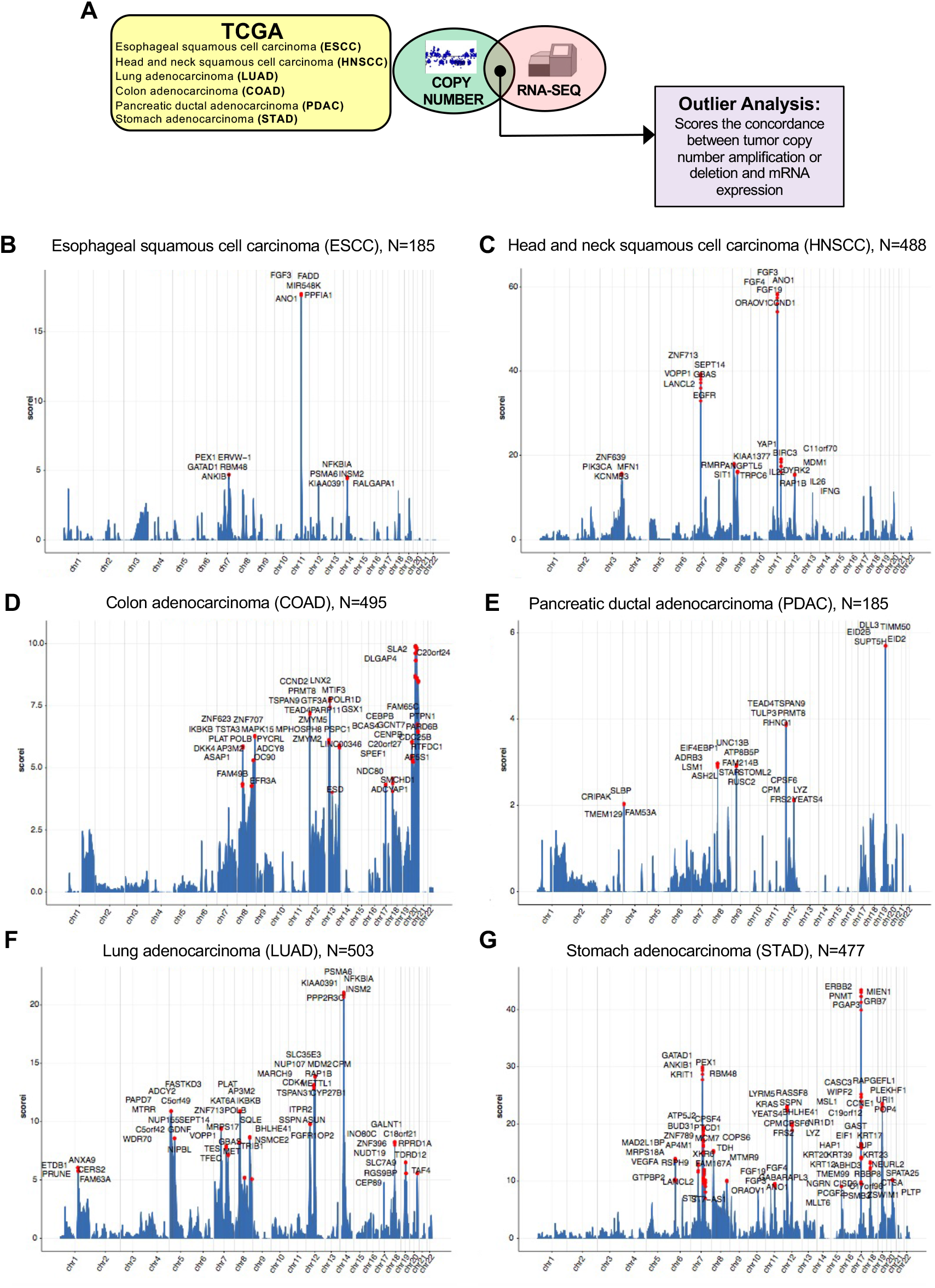
Overview of integrative analysis for expression and copy number amplification events as pan-cancer outliers. A) Schematic of integrative analysis to nominate outlier gene candidates. B) Genomic landscape of selected TCGA esophageal squamous cell carcinoma outliers. C) Genomic landscape of selected TCGA head and neck squamous cell carcinoma outliers. D) Genomic landscape of selected TCGA colon adenocarcinoma outliers. E) Genomic landscape of selected TCGA pancreatic ductal adenocarcinoma outliers. F) Genomic landscape of selected TCGA NSCLC adenocarcinoma outliers. G) Genomic landscape of selected TCGA stomach adenocarcinoma outliers.

### Derivation of a pan-cancer panel of first hit-engineered organoid models

We then evaluated the aforementioned cancer type-specific candidate overexpression outliers in minimally transformed organoid models containing single or double oncogenic mutations, generated from the corresponding tissues of origin. Primary epithelia isolated from human or mouse were cultured as previously described (Li et al., 2014; Lo et al., 2021; Neal et al., 2018; Salahudeen et al., 2020). We generated *Kras*^*G12D*^ mouse pancreatic organoids by culturing *LSL-Kras*^*G12D*^ (Hingorani et al., 2003) pancreas as air-liquid interface organoids and infected with adenovirus-Cre-GFP to activate expression of latent *Kras*^*G12D*^ (Li et al., 2014) **(Fig. S2A-B)**. Human wild-type gastric and colon organoids were grown under standard submerged methods (Lo et al., 2021; Matano et al., 2015; Sato et al., 2011). *APC*^*-/-*^ colon organoids were generated by CRISPR/Cas9 (Drost et al., 2015; Matano et al., 2015), while *TP53*^*R175H*^ gastric organoids were generated by stable lentivirus transduction **(Fig. S2C-F)**.

For this study, we also generated three novel organoid models corresponding to *p53*-null (p53^-/-^) oral squamous cell carcinoma, p53^-/-^ esophageal squamous cell carcinoma and *Kras*^*G12D*^; p53^-/-^ lung adenocarcinoma, described below. Mutational inactivation of the *p53* tumor suppressor gene occurs at high frequency of primary HPV-negative HNSCCs. We established normal oral mucosa (OM) organoids from mouse glossal epithelium cultured as air-liquid interface (ALI) organoids (**Fig. 2A**). Normal oral mucosal tissue consists of stratified squamous epithelium and connective tissue. In the basal layer, KI67, p63 and KRT5 are expressed (Ebrahimi and Botelho, 2017; Jones et al., 2019). Mouse OM ALI organoids recapitulated squamous epithelial structures (**Fig. 2A**). The basal cell marker KRT5 was expressed in the outer periphery, in which KI67+ cells were also detected, suggesting that cell proliferation occurred mainly at these sites (**Fig. 2A)**. These mouse OM organoids were maintained in ALI for more than 1 year.

**Figure 2.**
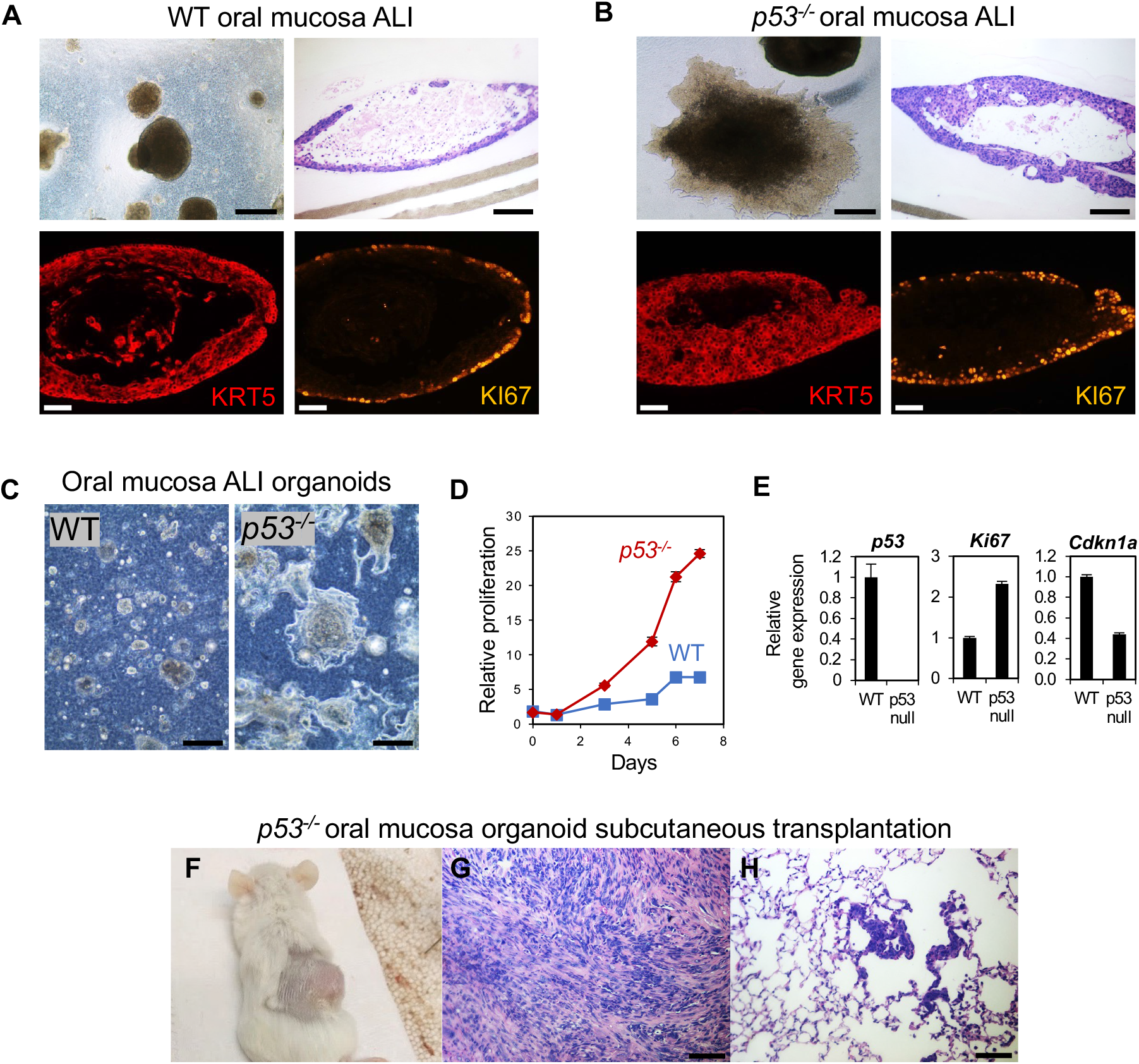
Generation and characterization of *p53*^*-/-*^ mouse oral mucosa organoids. A-B) WT (A) and *p53*^*-/-*^ mouse oral mucosa organoids (B) cultured in ALI for 34 days. Scale bar: brightfield; 500 μm (left top panel), H&E staining; 100 μm (right top panel), IF staining; 50 μm (bottom panels). C) Organoid morphology of WT (top panel) and *p53*^*-/-*^ mouse oral mucosa organoids (bottom panel) cultured in ALI for 7 days. Scale bar: 200 μm. D) Proliferation of oral mucosa *p53*^*-/-*^ and WT organoids in ALI culture as assessed by resazurin reduction. E) Expression of *p53, Ki67* and *Cdkn1a*, as assessed by RT-qPCR, in *p53*^*-/-*^ versus WT mouse oral mucosa organoids in ALI. F) Tumor formation of *p53*^*-/-*^ mouse oral mucosa organoids 7 months after implantation (WT; n=3, *p53*^*-/-*^; n=3). G) Primary tumor of p53^*-/-*^ mouse oral mucosa organoids from F, scale bar: 100 μm. H) Lung metastasis of p53^*-/-*^ mouse oral mucosa organoids from F, scale bar: 100 μm.

To initiate oncogenic transformation in vitro, mouse *p53*^*flox/flox*^ mouse OM organoids were infected with adenovirus Cre-GFP to create a contextual *p53*^-/-^ oral cancer model (**Fig. 2B-F**). These *p53*^-/-^ organoids exhibited a stratified cell arrangement with a thick KRT5+ cell layer and multilayered KI67+ cells upon long term culture **(Fig. 2B)**. *p53* deletion enhanced proliferation (**Fig. 2C,D**), accompanied by *Ki67* upregulation and *Cdkn1a* downregulation (**Fig. 2E**). Furthermore, *p53* deletion induced in vivo tumorigenicity and metastasis of mouse OM organoids **(Fig. 2F-H)**.

Mouse OM organoids could also be grown in submerged formats where they exhibited similar organization as ALI OM organoids with an outer rim of KRT5+ KI67+ proliferative basal cells and could again be cultured for more than 1 year (**Fig. S3A**). Human oral mucosa could be cultured for approximately 6 weeks in submerged or collagen air-liquid interface formats and included KRT5+ ECAD+ basal cells and the notable presence of keratinization within the organoid lumens (**Fig. S3B**,**C**). Upon lentiviral transduction with the oncogenic R175H allele of *p53*, these could be serially passaged up to 6 months but without obvious signs of dysplasia (data not shown).

In addition to oral squamous cell carcinoma, we also generated ALI esophageal squamous organoids using the same *p53*^*flox/flox*^ murine model (**Fig. 3A-C**). Esophageal organoids from *p53*^*flox/flox*^ mice formed KRT5+ squamous epithelium with similar morphology to OM organoids (**Fig. 3C**). Upon in vitro infection with adenovirus Cre-GFP, the resultant *p53*^*-/-*^ esophageal organoids exhibited dysplasia (**Fig. 3D**), maintenance of KRT5 (**Fig. 3E**), and loss of *p53* expression (**Fig. 3F**). Further, we observed in vivo tumorigenicity when *p53*^*-/-*^ ESCC organoids were subcutaneously transplanted into immunodeficient mice with keratin pearl formation and other histologic features consistent with squamous cell carcinoma (**Fig. 3G,H**).

**Figure 3.**
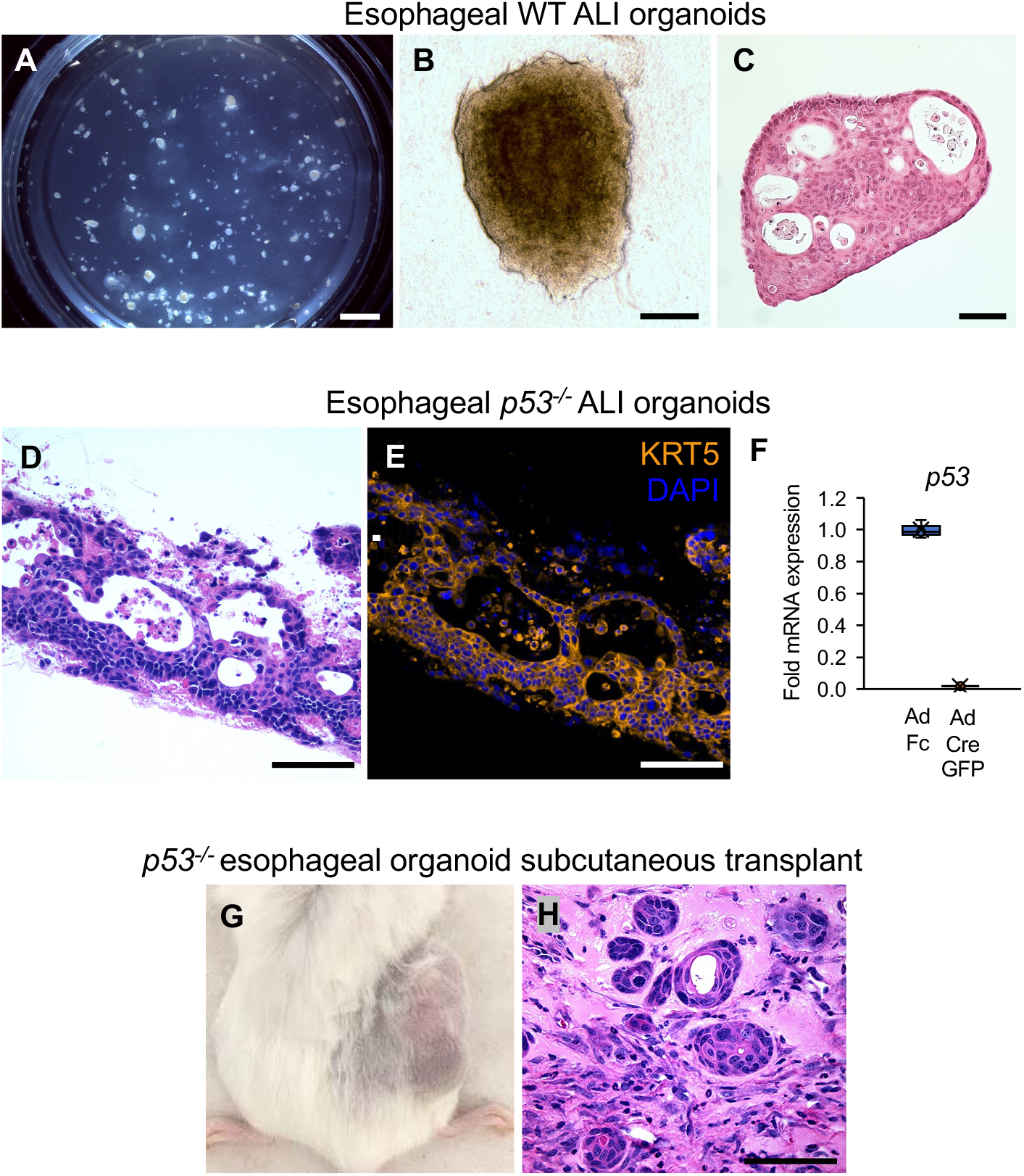
Generation and characterization of *p53*^*-/-*^ mouse esophageal squamous cell carcinoma organoids. A) Stereomicroscope images of wild-type esophageal organoids. Scale bar = 2.5 mm. B) Brightfield of wild type esophageal organoids. Scale bar = 250 um. C) H&E staining of wild-type esophageal organoids. Scale bar = 250 micron. D) H&E staining of p53^*-/-*^ esophageal organoids. Scale bar = 250 micron. E) KRT5 Immunofluorescence of (D). F) qRT-PCR of *p53* in Ad-Fc or Ad-Cre GFP infected *p53*^*flox/flox*^ esophageal organoids. G) Subcutaneous tumor formation 6 months post-implantation of *p53*^*-/-*^ esophageal organoids. H) H&E stain of (G), scale bar = 250 um.

Lastly, NSCLC lung adenocarcinoma organoids were generated from murine pulmonary parenchymal tissue using a protocol similar to our previous studies (Li et al., 2014; Salahudeen et al., 2020). Accordingly, ALI lung organoids were generated from unexcised LSL-*Kras*^*G12D*^; *p53*^*flox/flox*^ mice (Hingorani et al., 2003; Jackson et al., 2005). Upon infection with a negative control adenovirus expressing an immunoglobulin Fc fragment (Ad-Fc), these wild-type lung organoids proliferated in media containing EGF and Noggin (EN), but not in basal F12 media, similar to our prior studies in human lung organoids (Salahudeen et al., 2020). However, upon in vitro adenovirus Cre-GFP infection, the lung ALI organoids were converted to a *Kras*^*G12D*^; *p53*^*-/-*^ genotype and exhibited EGF independence (**Fig. 4A**), expression of mutant *Kras*^*G12D*^ and loss of *p53* (**Fig. 4B,C**), retention of TTF-1 expression (**Fig. 4D**), and in vivo tumorigenicity and metastases (**Fig. 4E-G**).

**Figure 4.**
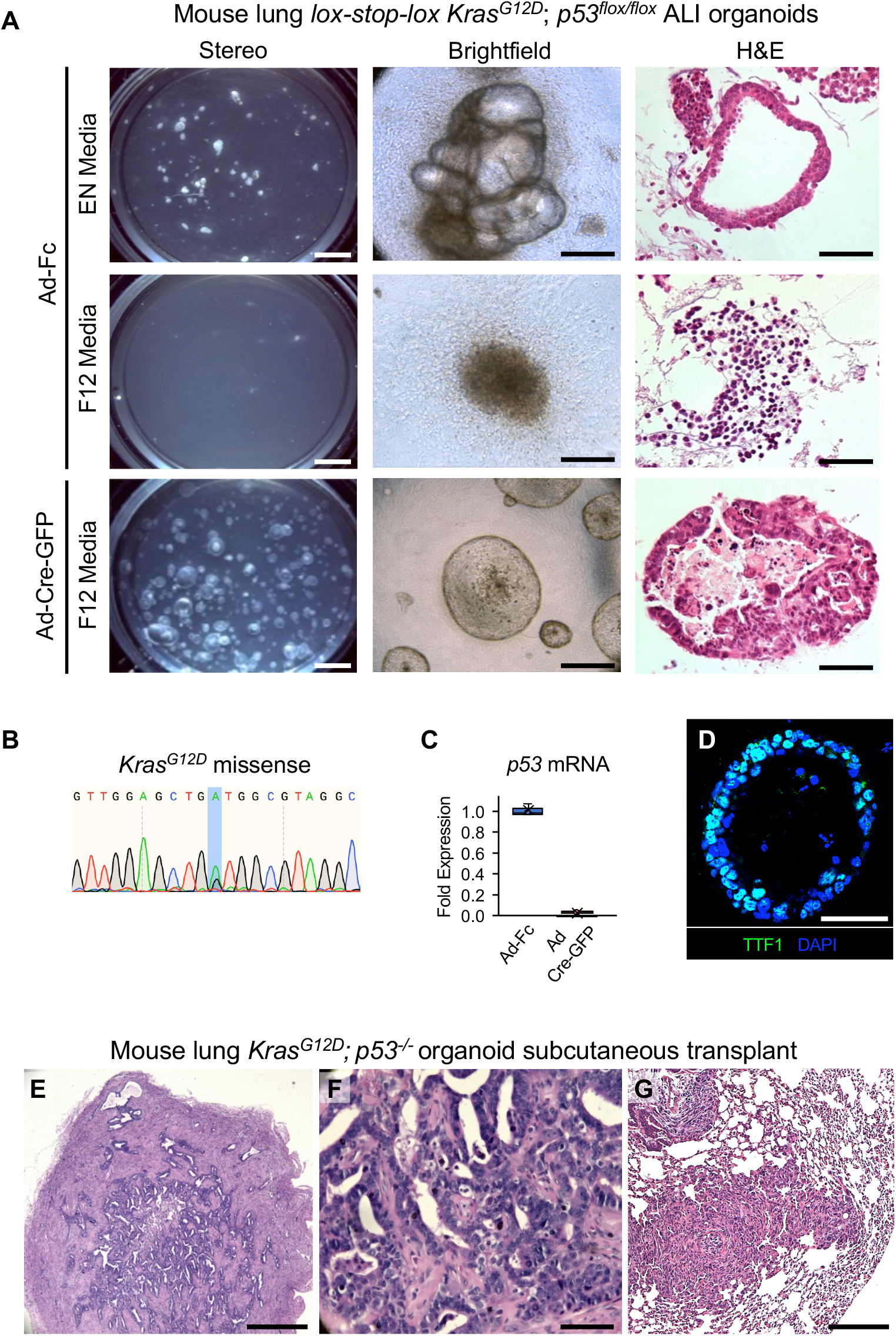
Generation and characterization of *Kras*^*G12D*^ *p53*^*-/-*^ lung adenocarcinoma organoids. A) Microscopy of *Kras*^*G12D*^; *p53*^*-/-*^ lung organoids with and without activation of latent alleles. Left, Stereomicroscopy of organoids at d28, Middle, phase contrast microscopy of organoids at d28, Right, H&E staining of organoid cultures at day 28. Scale bars = 3 mm, 250 µm, and 250 µm respectively. B) Sanger sequencing of *Kras* cDNA upon activation of latent LSL *Kras*^*G12D*^ from (A). C) qRT-PCR of *p53* in Ad-Fc or Ad-Cre GFP infected *Kras*^*G12D*^, *p53*^*-/-*^ lung organoids. D) TTF-1 immunofluorescence of *Kras*^*G12D*^, *p53*^*-/-*^ lung organoids, scale bar = 250 microns. E-G) H&E of primary tumor, high magnification, and lung metastasis 8 weeks post-s.c. implantation of *Kras*^*G12D*^, *p53*^*-/-*^ lung organoids. Scale bars = 500, 100 and 500 microns, respectively.

### Contextual tissue-specific functional screening of pan-cancer copy number amplification outliers

Having characterized these 6 tissue-specific, minimally-transformed oncogenic models, we then performed integrative analysis on the expression outliers for each cognate cancer type **(Fig. 1)** to computationally identify over 1000 candidate amplification outliers (**Table S1**). From these loci, we selected 393 available full-length cDNAs from the CCSB-Broad lentivirus ORF collection (Yang et al., 2011) having barcodes external to the ORFs, facilitating pooled screens. A custom ORF library was thus generated for each of the 6 cancer-specific outlier predictions, for infection of the corresponding minimally-transformed tissue organoids. For instance, esophageal ESCC TCGA outliers were represented by an equivalent barcoded lentivirus ORF library, for infection into p53^-/-^ esophageal organoids **(Fig. S1B, Fig. 5)**.

**Figure 5.**
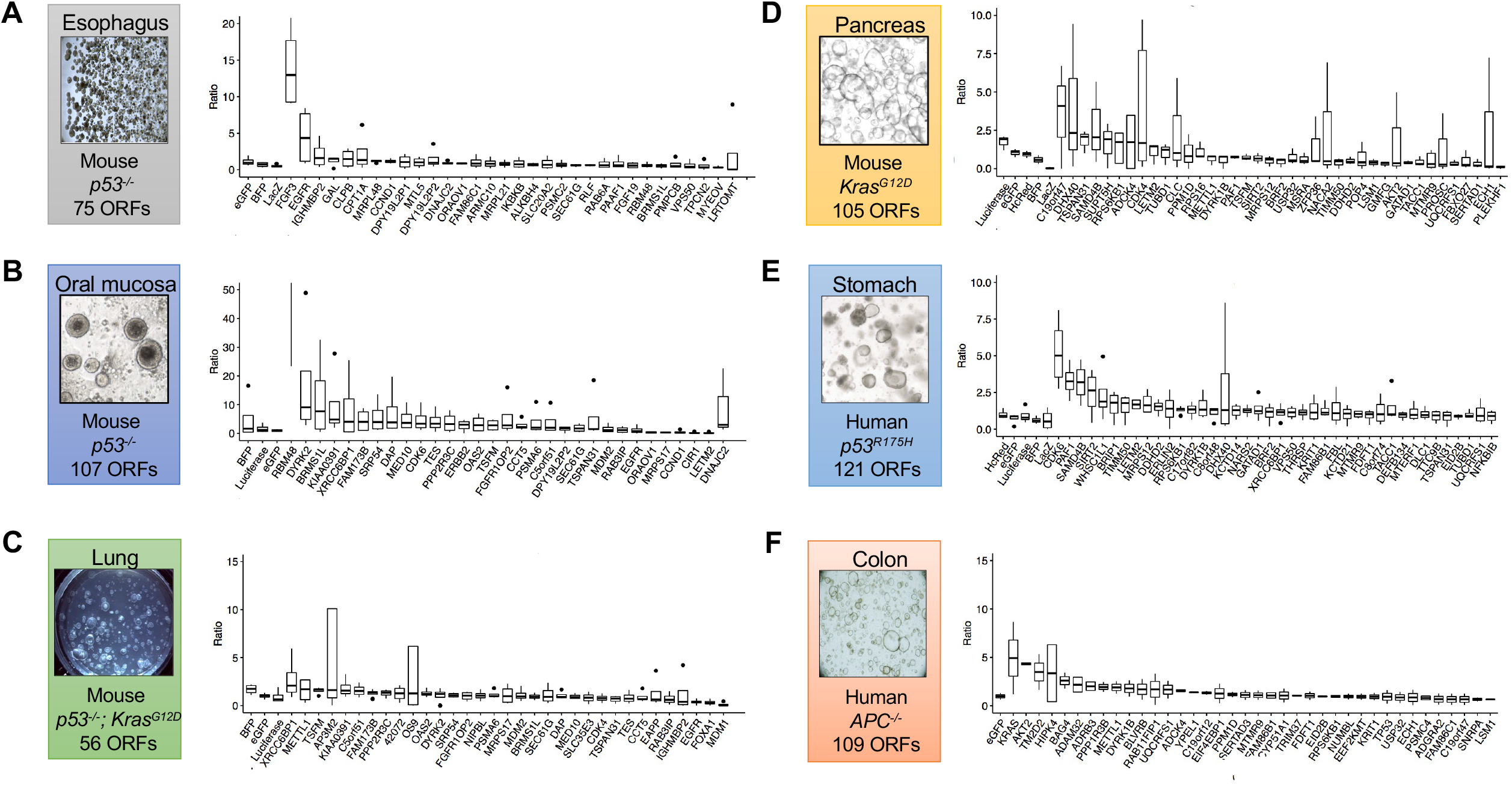
Screening pan-cancer candidate amplification outliers in organoids with corresponding tissue context. A-F) Barcode ratios of terminal:initial timepoints demonstrating relative enrichments from pooled lentiviral ORF screens. Boxplots represent four technical replicates with the exception of F) which was performed with two technical replicates.

Lentiviruses were generated in arrayed format and pooled for equal titer amounts **(STAR Methods)**. Loci encoding already-established oncogenic drivers were deliberately not included in our screen to avoid potentially dominant “jackpot” effects that could obscure contributions of novel outliers. Organoids were infected with the pooled virus corresponding to the outliers for that histologic site and screened as independent technical replicates, and aliquots from initial plating (t=0) and after four passages (terminal time point) were collected. NGS sequencing was performed to evaluate barcode enrichment at the terminal time point, typically at culture day 30-50 days, versus t=0 (**Fig. 5A-F**). Organoids were subjected to prolonged culture after pooled lentiviral infection and puromycin selection; barcodes were quantitated at each time point by NGS. Barcode distribution consistency was observed across technical replicate screens, typically n=4 (**Fig. S4-6**). Notably, outliers were functionally enriched in cell cycle processes (CDK4, CDK6 in gastric and pancreas), DNA repair (XRCC6BP1 in lung), and kinase signaling (KRAS, AKT in colon) (**Fig. 5A-F**). Evaluation of these genes and other enriched outliers will be undertaken in these adenocarcinoma models in future work. Interestingly, in *p53*^*R175H*^ human gastric organoids, two of the top five outliers corresponded to *CDK6*. When analyzing gastric cancer outliers that displayed >2-fold change vs. t=0, six of the top hits (DYRK1B, SAMD4B, MRPS12, SIRT2, EIF3K, and PAF1) all co-localized to chromosome 19q13.2.

Screens in the *p53*^-/-^ oral mucosa organoid model identified the Dual Specificity Tyrosine Phosphorylation-regulated kinase 2 (DYRK2) as one of the most highly-ranked hits (**Fig. 5B**). *DYRK2* amplification is seen in approximately 5% of HNSCC (Cancer Genome Atlas, 2015). *DYRK2* is thought to be required for tumor growth via proteasome phosphorylation (Guo et al., 2016) and induces p53-dependent apoptosis to DNA damage (Taira et al., 2007; Tandon et al., 2021). Of the *DYRK2* ORFs screened, we found greater enrichment of ORFs mapping to the isoform 2 transcript variant. To iteratively validate the role of DYRK2, we infected *p53*^*-/-*^ mouse oral mucosa organoids with DYRK2 isoform 2-expressing lentivirus (**Fig. 6A**). DYRK2 overexpression enhanced proliferation of *p53*^*-/-*^ oral mucosa organoids versus lentivirus eGFP-infected organoids in vitro (**Fig. 6B,C)** and promoted *in vivo* tumorigenicity upon subcutaneous transplantation (**Fig. 6D,E**).

**Figure 6.**
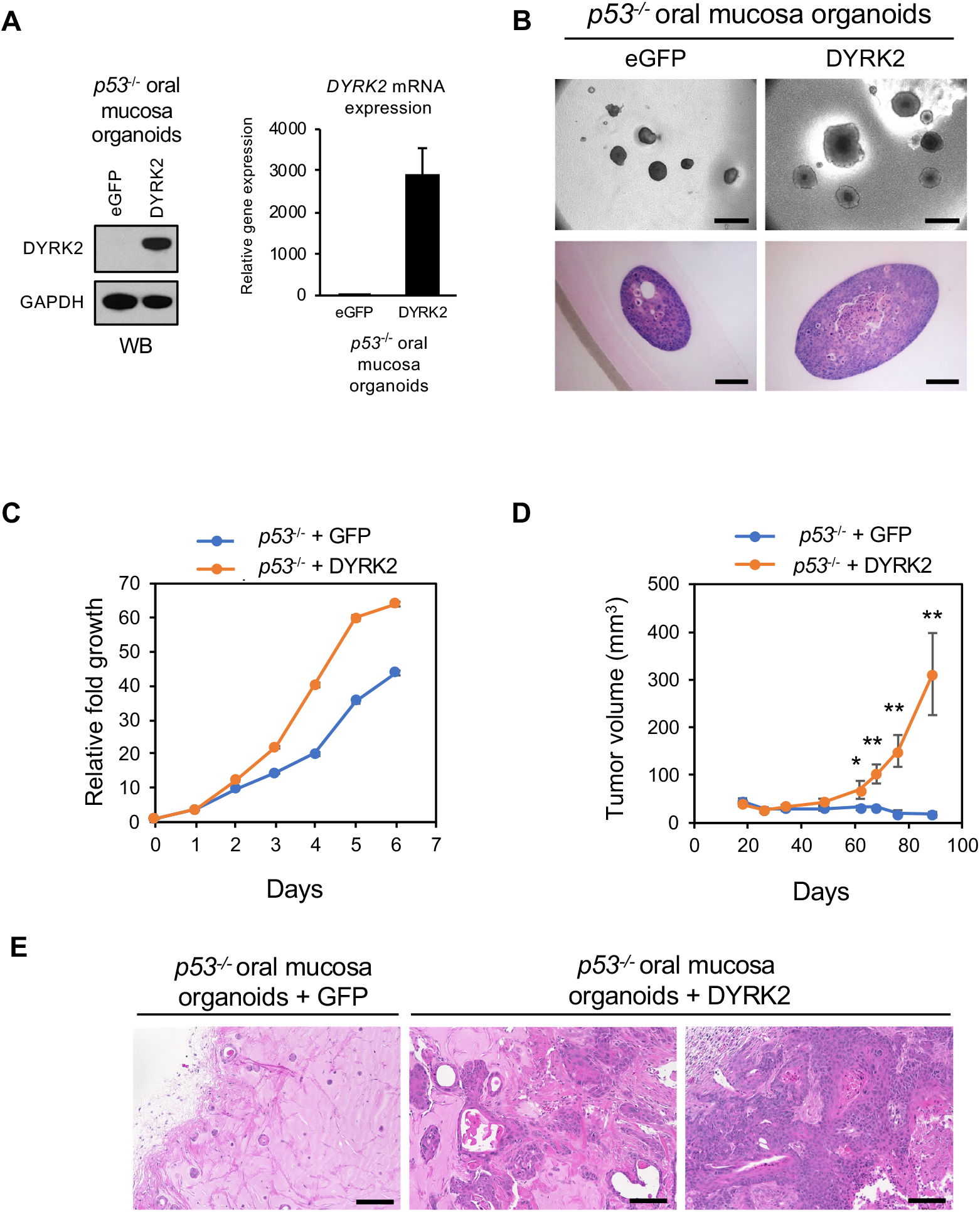
DYRK2 overexpression induces proliferation and tumorigenicity of *p53*^*-/-*^ mouse oral mucosa organoids. A) Validation of DYRK2 overexpression by immunoblotting (left panel) and qRT-PCR (right panel). B) Morphology of *p53*^*-/-*^ eGFP and *p53*^*-/-*^ DYRK2 organoids cultured in ALI for 14 days. Scale bar: bright field; 500 μm (top panel), H&E staining; 100 μm (bottom panel). C) Proliferation of *p53*^*-/-*^ eGFP and *p53*^*-/-*^ DYRK2 in ALI as assessed by resazurin reduction. D) Tumor volume of *p53*^*-/-*^ eGFP and *p53*^*-/-*^ DYRK2 (*p53*^*-/-*^ eGFP; n=6, *p53*^*-/-*^ DYRK2; n=7). Data represent mean ± SEM. Two-sided P-values were calculated by Student’s t-test. *p < 0.05, ** p < 0.01. E) H&E staining of *p53*^*-/-*^ eGFP and *p53*^*-/-*^ DYRK2 tumors. Scale bar: 100 μm.

### FGF3 is a candidate amplified oncogenic driver in esophageal squamous cell carcinoma

Results from our lentivirus pooled ORF expression screen also identified *FGF3* and *EGFR* enrichment in *p53*^*-/-*^ ESCC organoids with *FGF3* having the greatest overall effect in this system; a comparatively modest enrichment was also observed for *CCND1* (**Fig. 5A**). FGF3 was lentivirally expressed in an independent ESCC *p53*^*-/-*^ organoid line not used in the ORF screen (**Fig. S7A**,**B**), again confirming increased proliferation in serum-containing minimal F12 medium (**Fig. 7A-C**). Given the strong proliferation phenotype in the absence of the mitogen EGF, we hypothesized that FGF3 overexpression could serve an EGF surrogate, and that FGF3 could act functionally as an autocrine growth factor in ESCC. Such a model was supported by our finding that FGF3 was indeed secreted into culture supernatants as confirmed by ELISA (**Fig. S7B**).

**Figure 7.**
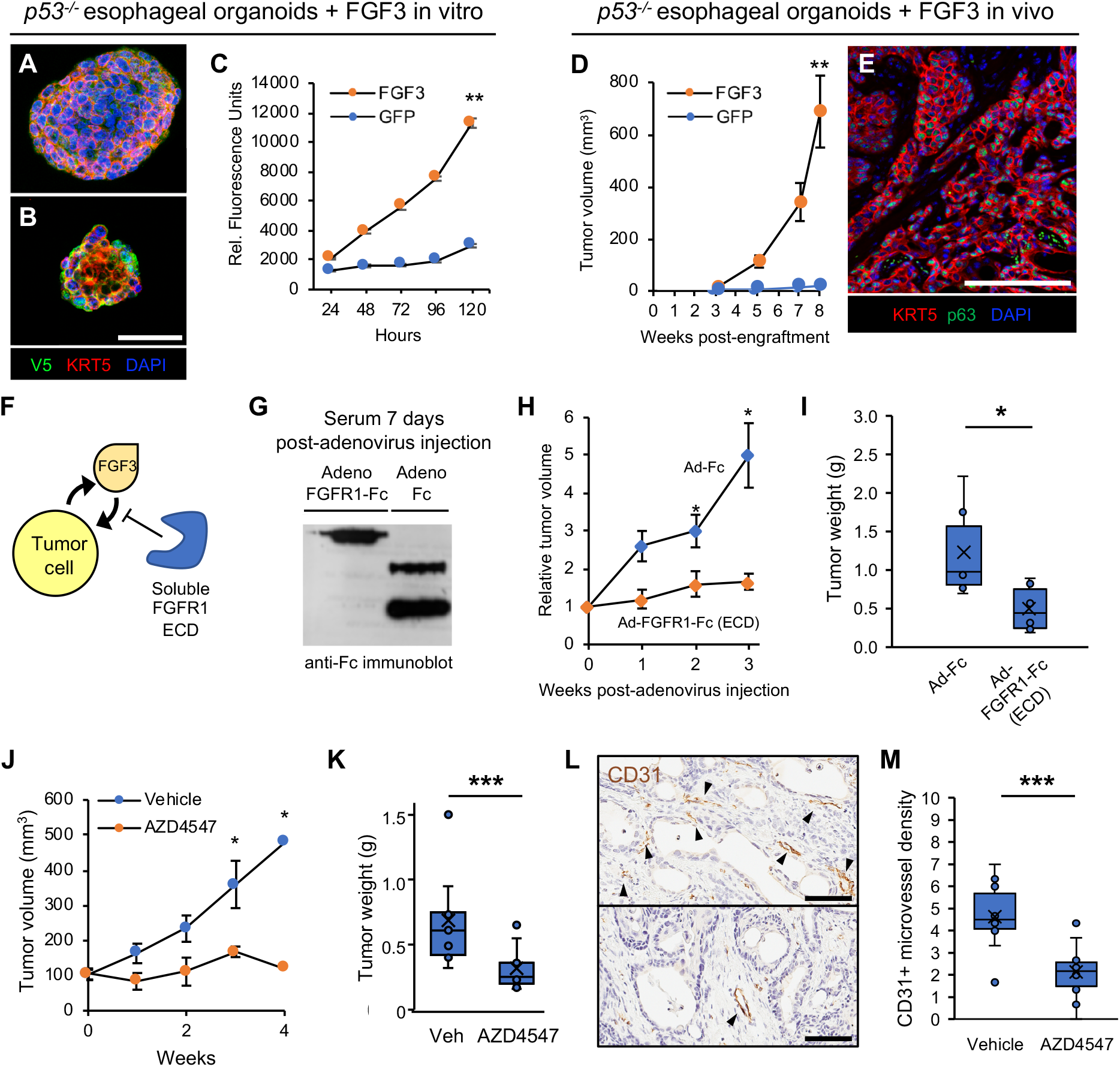
Iterative validation of FGF3 amplification as an oncogenic driver in *p53*^*-/-*^ mouse esophageal squamous organoids. A) Multicolor immunofluorescence of p53^-/-^ esophageal organoids with lentiviral expression of FGF3 with a C-terminal V5 tag. B) Multicolor immunofluorescence of p53^-/-^ esophageal organoids with lentiviral expression of GFP with a C-terminal V5 tag. C) in vitro proliferation of FGF3 versus GFP-expressing *p53*^*-/-*^ esophageal organoids by resazurin reduction. D) in vivo tumor formation and growth upon subcutaneous transplantation of FGF3-versus GFP-expressing p53^-/-^ esophageal organoids in immunodeficient NOG mice. Each group has n = 10 biological replicates. E) Multicolor immunofluorescence of FGF3 p53^-/-^ esophageal organoid subcutaneous tumor sections. F) Schematic of hypothesized autocrine mechanism of FGF3 driven oncogenesis and potential disruption by a soluble ligand-binding extracellular domain of FGFR1. G) Anti-Fc immunoblot showing serum expression of FGFR1-Fc or Fc alone at 2 days after i.v. injection of immunodeficient NSG mice infected with the corresponding recombinant adenoviruses. H) Serial measurements of subcutaneous tumor growth subsequent to adenoviral infection of FGFR1 ECD-Fc versus Fc controls in FGF3 expressing p53^-/-^ esophageal organoid subcutaneous tumors. Each group has n = 6 biological replicates. I) Terminal tumor weights of H, *P =* 0.026. J) Serial measurements of subcutaneous tumor growth subsequent to daily administration of vehicle or the FGFR inhibitor AZD4547. Each group has n = 10 biological replicates. K) Terminal tumor weights of J, *P* = 0.00057. L) Representative CD31 immunohistochemistry staining of subcutaneous tumors from J. Scale bar = 200 µm. M) Chalkley quantitation of CD31+ microvessel density in (L), *P =* 0.001. Each group had three technical replicates for each biological replicate.

We then investigated if FGF3 overexpression conferred a phenotype upon subcutaneous transplantation into immunodeficient mice. Paralleling in vitro observations, FGF3 overexpression in *p53*^*-/-*^ esophageal organoids induced robust tumorigenicity in vivo **(Fig. 7D-E; Fig. S7C)**. We then assessed if tumor growth could be attenuated by FGFR antagonism via expression of a soluble ligand-binding FGFR1 ectodomain (ECD) fused to an antibody Fc fragment (FGFR1-Fc) to scavenge secreted FGF3 (**Fig. 7F**). To this end, we utilized an adenoviral vector (Ad-FGFR1-Fc) for in vivo liver infection and hepatocyte secretion of the FGFR1-Fc ectodomain fusion protein into the circulation of mice (**Fig. 7G**). Mice infected with FGFR1-Fc adenovirus but not a control adenovirus expressing an Fc immunoglobulin fragment alone (Ad-Fc) inhibited tumor growth (**Fig. 7H-I**) in FGF3-overexpressing organoid tumors.

Having established tumor suppression by circulating FGFR1-Fc, we then evaluated whether selective small molecule FGFR tyrosine kinase inhibitors, a drug class efficacious in treatment of *FGFR*-rearranged tumors (Weaver and Bossaer, 2021), could also inhibit proliferation of ESCC overexpressing *FGF3*. We employed two first generation pan-FGFR inhibitors, AZD4547 and BGJ-398 and observed a clear dose dependent inhibition of *p53*^-/-^ ESCC organoid models overexpressing FGF3 with nanomolar EC_50_ values (**Fig. S7D**). Interestingly, this response to FGFR inhibitors was masked when organoids were cultured in EGF (data not shown) possibly due to functional redundancy between these two growth factor classes. Accordingly, AZD4547 significantly reduced growth of FGF3-overexpressing *p53*^*-/-*^ esophageal organoid tumors versus vehicle (**Fig. 7J-K**), Importantly, growth of GFP tumors (i.e. without FGF3 overexpression) was not significantly different between AZD4547 and vehicle treatments (**Fig. S7E-F**). Given the importance of FGF signaling in angiogenesis, we then evaluated whether FGFR inhibition resulted in decreased tumor vasculature. FGF3-overexpressing organoid tumors observed decreased CD31^+^ microvessel density upon AZD4547 treatment compared to vehicle (**Fig 7L,M**). Taken together, our findings support autocrine FGF-FGFR signaling as a potential druggable oncogenic mechanism in esophageal squamous tumors harboring *FGF3* amplification and overexpression.

## DISCUSSION

Pan-cancer bioinformatics studies of TCGA and other genome-scale cancer surveys have proven informative but lack a laboratory counterpart of functional validation in an equivalent in vitro contextual experimental system (Hahn et al., 2021). The advent of organoid technologies now powerfully enables systematic interrogation of these multi-cancer datasets in cognate pan-organoid screens matched to the tissue of origin. Here, we present a pan-cancer approach to evaluate the oncogenic potential of putative copy number alterations. First, an integrative analysis of gene expression with copy number alterations prioritized and nominated copy number outliers with potential significance in cancer biology. Second, these candidate SCNA datasets were directly coupled to functional validation in primary organoid cultures spanning colon, stomach and pancreas and three novel organoid models of oral mucosa, lung, and esophagus.

The present method utilized contextual modeling where SCNA outlier candidates specific to a cancer histologic type were screened in matching tissue organoids that contained a pre-existing, predominant “first hit” for that malignancy: for instance, *KRAS* mutations in PDAC, *APC* in COAD and *p53* mutation in HNSCC, ESCC, LUAD and STAD. This not only modeled the physiologic occurrence of SCNAs in the background of such pervasive signature mutations but also conveniently exploited the robust growth of such single oncogene-engineered organoids versus their wild-type counterparts to amass sufficient starting material for the barcoded screens. Further, the “bottom-up” nature of the current screens, against a minimally transformed background, conveyed potential advantages of increased sensitivity to detect oncogenic loci, versus functional testing in a patient-derived tumor organoids or cell lines where a multitude of pre-existing epi/genetic alterations could obscure effects of a given outlier candidate.

Copy number alterations are one of the most ubiquitous features of cancer genomes yet a comprehensive understanding of their oncogenic contributions has yet to be achieved (Kristensen et al., 2014). Earlier nominations of copy number events have primarily focused on delineating regions of loss or gain with high genomic resolution but have placed less emphasis on nominating regions of functional consequence due to concomitant changes in gene expression. While chromosomal deletions involving canonical tumor suppressors or amplifications of growth factor receptor tyrosine kinases or RAS kinases are well characterized, progress to evaluate other copy number alterations have been modest. Functional genomics experiments involving immortalized cancer cell lines have identified synthetic lethality events such as *MTAP* deletion and S-adenosyl methionine arginine methyltransferase (Kryukov et al., 2016) or a recent preprint suggesting inhibition of PKMYT1 in CCNE1 amplified tumors (Gallo et al. bioRxiv 2021). Future efforts involving panels of thousands of immortalized cell lines in large scale small molecule and functional genomic experiments such as the Cancer Dependency Map have the potential to yield further insights into these copy number alterations (Boehm et al., 2021; Hahn et al., 2021). Here, we prioritized screening of amplified outliers, reasoning that oncogenic hits would be amenable to pharmacologic inhibition, but the present approaches could be extended to CRISPR/sgRNA screens for deletion outliers.

We identified several amplification outliers with known oncogenic potential in models of colon, pancreatic, and gastric cancer. Here, the interrogation of 121 amplification outlier ORFs from the TCGA STAD dataset in *p53*^*R175H*^-overexpressing human gastric organoids revealed the highest scoring hit as CDK6, a commonly amplified gene widely implicated in carcinogenesis and a target for pharmacological inhibition. The ability of the CDK4/6 inhibitor palbociclib to induce senescence in gastric cancer cells is reduced upon *p53* knockdown, consistent with cooperation between *p53* mutation and *CDK6* in promoting tumorigenesis (Valenzuela et al., 2017). Intriguingly, several of the top gastric cancer SCNA hits, namely *DYRK1B, SAMD4B, MRPS12, SIRT2, EIF3K*, and *PAF1*, all co-localized to chromosome 19q13.2, suggesting a multifaceted role of this amplicon in the fitness of gastric cancer cells. As amplification of chromosome 19q13.2 occurs in other cancer types including pancreas (Kuuselo et al., 2010), lung (Kim et al., 2005), and breast (Basu and Lambring, 2021; Bellacosa et al., 1995), potential cooperation between these SCNA hits warrants further investigation.

We also recognized and functionally validated two novel oncogenic amplifications in squamous cancers of the oral cavity and esophagus. In contrast to adenocarcinomas, squamous cancers exhibit relatively fewer actionable mutations. Here, DYRK2 strongly promoted in vitro proliferation and in vivo tumorigenicity of oral mucosal organoids, as a highly relevant model for head and neck squamous cell carcinoma. DYRK2 is a family member of relatively understudied tyrosine kinases that phosphorylate histones and other substrates (Tandon et al., 2021). Our studies suggest the potential dependency of DYRK2-amplifications in oral cancers and corresponding efficacy of DYRK2 selective inhibitors.

The functional validation of *FGF3* as an amplified oncogene at the 11q13 locus using *p53*^*-/-*^ esophageal organoids directly suggests that this growth factor could represent an autocrine vulnerability in amplified tumors, FGFR small molecule inhibitors have recently received FDA approval in biliary and bladder cancer (Weaver and Bossaer, 2021). Our preliminary findings of in vivo and in vitro efficacy of FGFR inhibitors against FGF3-overexpressing esophageal organoids warrant further exploration and evaluation in FGF3-amplified patient derived models of ESCC as well as and other cancer types such as the copy number-driven integrative breast cancer subgroup, IntClust2, which manifests prominent FGF3 amplification (Rueda et al., 2019).

Notably, *EGFR* and *KRAS* amplifications confer sensitivity to inhibitors targeting corresponding pathways in cancer patients (Catenacci et al., 2021; Wong et al., 2018). In addition, FGF3 is an embryonal FGF and is not significantly expressed in adult human tissues; thus FGF3 could be considered an oncofetal antigen susceptible to adoptive T cell therapies selective for cells presenting FGF3 peptides on class I MHC. Given that more than 40% of ESCC harbor FGF3 amplifications within the 11q13 amplicon (Cancer Genome Atlas Research et al., 2017), successful targeting of FGF3 in ESCC could have significant clinical impact (Abnet et al., 2018). However, whether these patients may respond to FGFR inhibition requires further study, as other likely driver loci such as *CCND1* are co-amplified with *FGF3* at 11q13, although in our screen *FGF3* displayed stronger effects than *CCND1*. Nevertheless, our data identify *FGF3* as an additional driver at this locus which could indeed cooperate with *CCND1*.

Overall, we have demonstrated that primary organoid culture can be robustly applied to validate pan-cancer genomic datasets in pooled barcoded screening formats, using amplified outlier loci as proof-of-principle. However, there are several limitations to the present study. First, the usage of cDNA ORFs in lentiviral vectors is subject to inherent packaging limits where candidate genes associated with large inserts may not be feasibly generated with sufficient titer. Furthermore, achieving uniform library representation via tittering and pooling arrayed lentiviral preps may be cumbersome and costly. Desired full-length cDNAs either may not be available or may not represent the correct isoform in the tissue specific context of a given organoid model, precluding evaluation of given loci. Future efforts could adapt this pan-cancer organoid framework to utilize CRISPR activation or CRISPRon where dCas9 is fused to transcriptional activators such as VPR or to histone demethylases (Gilbert et al., 2014; Horlbeck et al., 2016; Nunez et al., 2021). Lastly, the use of exclusively human organoid models and progressively miniaturized formats (Du et al., 2020) could be combined with systematic functional evaluation of additional tissue organoid types, genomic regions and classes of genetic alterations.

## ACKNOWLEDGEMENTS

We thank members of the Kuo laboratory and the CTD^2 consortium for helpful discussions. We thank Scott Younger for supervising the design and construction of lentiviral library pools. This work was supported by the NCI Cancer Target Discovery and Development (CTD2) Network (U01CA217851, C.J.K and C.C.; U01CA176058, W.C.H.), Support was also provided by NIH U54CA224081, NIH U01CA199241, Emerson Collective, Ludwig Cancer Research and Stand Up To Cancer to C.J.K. This manuscript is dedicated to the memories of Dr. Daniela Gerhard and Dr. Kenneth Scott.

## STAR Methods

### Lead Contact and Resource Availability

Further information and requests for resources and reagents should be directed to and will be fulfilled by the Lead Contact, Calvin J. Kuo.

### Data and Code Availability

Code for Outlier Analysis and screening analysis will be made available in a Github repository upon publication of this manuscript.

Outlier screening data will be deposited at the CTD^2 portal upon publication of this manuscript: (https://ocg.cancer.gov/programs/ctd2/data-portal)

## EXPERIMENTAL MODEL AND SUBJECT DETAILS

### Human specimens

Normal tissues were obtained through the Stanford Tissue Bank from patients undergoing surgical resection at Stanford Health Care (SHC) All experiments utilizing human material were approved by the SHC Institutional Review Board and performed under protocols #28908. Written informed consent for research was obtained from donors prior to tissue acquisition. Analysis of influence of gender identity upon experiments was not performed.

### Mouse Models

Female C57BL/6 mice or NOG-E mice were used for organoid model generation and subcutaneous tumor implantation (mice were obtained from Taconic Biosciences) in accordance with NIH and Stanford Administrative Panel on Laboratory animal Care (APLAC). Mice were housed in pairs and used for experimentation at 4-8 weeks of age. Animals were maintained on a 12-hour light/dark cycle, in a temperature- and humidity-controlled room with food and water.

## METHOD DETAILS

### Computational amplified/upregulated outlier analysis

TCGA expression data (rsem genes normalized rnaseqv2) and copy number data (hg19 nocnv) were downloaded for lung adenocarcinoma (LUAD), esophageal carcinoma (ESCA, filtered to retain only the squamous samples), head and neck squamous carcinoma (HNSC), pancreatic adenocarcinoma (PAAD), colon adenocarcinoma (COAD) and stomach adenocarcinoma (STAD) from Firehose (http://gdac.broadinstitute.org/). We considered a segment amplified if the segment value was higher than the median value of all segments of the sample plus six times the standard deviation of the central quantiles of the values of the sample (0.25-0.75). Amplified segments were matched with the position of the gene (hg19) to assign gene-level values. For gene expression data, values were normalized using a log_2_ (adding a unit to the original count) transformation. We generated, for each gene a theoretical normal distribution, and obtained the 5% and 95% threshold quantiles where samples with real values higher than the 95% were considered *upregulated* expression outliers. Finally, we called a given gene an outlier if was simultaneously *amplified* and an *upregulated* expression outlier following the approach from (Curtis et al. Nature 2012). Outlier distributions were plotted with the ggbio package, transitions across genes were smoothed using the smooth.spline function.

### Organoid derivation and culture

Mouse esophagus, tongue/oral mucosa and pancreas were dissected from 6–8-week-old *p53*^*flox/flox*^ mice and mouse peripheral lung from 6–8-week-old *LSL-Kras*^*G12D*^; *p53*^*flox/flox*^ mice (Hingorani et al., 2003; Jackson et al., 2005) and minced into small pieces. Minced tissues were embedded in collagen gel in air-liquid interface (ALI) (Li et al., 2014; Nadauld et al., 2014). 2×10^8^ pfu adenovirus Ad-Fc or Ad-Cre-GFP (University of Iowa Vector Core) per 500 µl medium were added on top of the inner dish collagen I (Wako) to activate *Kras*^*G12D*^ expression and delete *p53*. Established organoids were maintained in ALI. Culture medium for mouse oral mucosa, esophagus and lung organoids was advanced DMEM/F12 supplemented with 1 mM HEPES, 10 mM nicotinamide, Glutamax, 1 mM N-acetylcysteine, B27, 0.5 μM A83-01, PSQ, 50 ng/ml recombinant human EGF, 100 ng/ml recombinant human NOGGIN. Pancreas organoids were cultured in WENR media (see below).

Human gastric and colon organoids were generated from deidentified surgical specimens from Stanford Hospital under an approved IRB protocol following established methods and grown in a submerged format in WENR media (Matano et al., 2015; Sato et al., 2011) in BME-2 extracellular matrix (R&D Systems) (Lo et al., 2021; Salahudeen et al., 2020).

The R175H mutant of p53 was transduced by lentivirus into wild type human gastric organoids APC null colon organoids were generated as previously described (Lo et al., 2021; Matano et al., 2015; Sato et al., 2011). For DYRK2 validation experiments, lentiviral DYRK2 (TRCN489007, Broad ORF) was infected into p53-null mouse oral mucosa organoids in log phase using spinfection (Lo et al., 2021; Salahudeen et al., 2020).

### Tissue contextual pooled screening of putative outlier genes

Organoids were either removed from matrix with either TrypLE or collagenase type IV as previously described (Lo et al., 2021; Neal et al., 2018). Upon matrix removal, organoids were further digested into single cell suspensions with TrypLE for 20 minutes at 37 degrees Celsius. Cells were centrifuged at 600 x g for 3 minutes, then washed and incubated with 100 Kunitz Units DNase I in 1 ml of Advanced DMEM/F12 for 15 minutes at room temperature. Cells were then counted and 1000 cells per ORF construct were infected via spinfection as four technical replicates. Each technical replicate was resuspended in complete culture media + 8 µg/ml polybrene, plus pooled lentivirus library with equal weighting inclusive of negative controls (MOI = 0.8) to a total volume of 250 µL per replicate in a 48 well plate. Plates were centrifuged for 1 hour at 32 degrees Celsius at 600 x g, then allowed to recover for 4-6 hours at 37 degrees in a cell incubator. Spinoculated cells were then plated in ECM/Matrigel at 100,000 cells per 50 µl droplet in complete media. Each technical replicate was cultured separately including subsequent passaging and gDNA harvesting. Each replicate was allowed to grow for 96 hours and then transferred either to ALI or maintained in ECM per above culture conditions. After 96 hours, cultures were then subjected to puromycin selection at IC_90_ concentrations established by prior dose response studies. After 96 hours of puromycin selection, the cultures were digested into single cell suspensions and half the biomass was snap frozen as a representation of the initial screen timepoint. Cultures were then passaged serially upon confluence as previously described (Lo et al., 2021; Neal et al., 2018) and screens were terminated after the fourth passage.

### Barcode Amplification and Deep Sequencing

Snap frozen cell pellets were extracted with DNeasy (Qiagen) following the manufacturer’s protocol. 10μg of genomic DNA was subjected to a one step PCR strategy per the Broad Genomic Perturbation Platform protocol (https://portals.broadinstitute.org/gpp/public/resources/protocols). Briefly, a pool of Illumina P5 primers were pooled and combined with a barcode library specific library primer. PCR products were gel extracted and subjected to Bioanalyzer analysis to assess for sample purity and quantified by Qubit. Samples were then pooled and Deep Sequencing was performed on an Illumina MiSeq instrument configured 300 | 8 | 0 | 300 bp with Nextera XT format.

Fastq files were processed using the R package ShortRead (Morgan et al., 2009) and ORF barcodes were counted as a DNAStringSet from the BioString R package (H. Pagès, 2021), and reverse barcodes were counted using the reverseComplement function. Barcodes (both forward and reverse) were counted using the vcountPattern function with an error limit of one single mismatch in forward and reverse strand reads combined.

### ORF enrichment analysis

The proportion of barcodes for each gene/sample were calculated by dividing the barcode counts by the total read counts of the NGS library. We then calculated the ratio of enrichment for each gene by dividing the proportion of barcodes at the terminal timepoint by the proportion of barcodes at the initial timepoint. In order to avoid division by zero, genes were filtered for any sample proportions less than 0.0003. Barcodes were plotted in rank order by genes with the median value of the ratio, separating the controls from the putative drivers.

### Immunoblotting

Lysate preparation and immunoblot analyses were performed using standard methods. Briefly, cells were harvested in RIPA buffer containing protease inhibitor cocktail (cOmplete Mini, Roche) and centrifuged at 5000 × g for 10 min to remove debris. Protein concentration was assessed using BCA Kit (Bio-Rad). Samples were supplemented with SDS sample buffer containing 5% 2-mercaptomethanol and 100 mM DTT. NuPAGE 4%–12% Bis-Tris Gels (ThermoFisher Scientific) were used for SDS-PAGE, then transferred to PVDF membranes (EMD Millipore). Membranes were blocked with 5% non-fat dry milk in TBS/0.05% Triton-X (TBST/milk). Membranes were incubated with primary antibodies: anti-DYRK2 antibody (#8143, Cell Signaling) and anti-GAPDH (#5174, Cell Signaling) diluted in TBST/milk overnight at 4°C. HRP-conjugated anti-Rabbit IgG (H+L) antibody (111-035-003, Jackson ImmunoResearch) were incubated for 1h at room temperature. Bound antibodies were visualized using SuperSignal West Pico Chemiluminescent Substrates (ThermoFisher Scientific) and exposure of AccuRay Blue X-Ray Films (E&K Scientific).

### qRT-PCR

Total RNA was isolated from cells using RNeasy (QIAGEN) and cDNA was synthesized using iScript Reverse Transcription Supermix (Bio-Rad). RT-qPCR was performed with Power SYBR Green assay (Applied Biosystems). Relative RNA expression was calculated using standard curve method and normalized by *Gapdh*.

### Proliferation assays and in vitro small molecule studies

Organoids were dissociated into single cells. 5000 cells were plated with 5-10 µl of Matrigel in flat-bottom 96 well plate or 10 µl of collagen in round-bottom 96 well plate. 10% AlamarBlue was added to wells and incubated for 4 hours at 37°C at indicated days. Fluorescence (Ex/Em=530/590 nm) was measured in a Biotek plate reader according to the manufacturer’s protocols. Small molecule treatments were carried out as dose response studies in triplicate 72 hours after seeding.

### Immunofluorescence staining

Paraffin-embedded sections were incubated in citrate antigen retrieval solution and blocked with 10% normal donkey serum. Sections were incubated with the following primary antibodies: mouse anti-E-cadherin (BD610182, BD Biosciences), Alexa Fluor 647-labeled anti-KRT5 (ab193895, Abcam), rat-anti-Ki67 (14-5698-82, eBioscience) overnight at 4°C. Sections were washed and subsequently incubated with the following secondary antibodies: Cy3-conjugated Affinipure goat anti-mouse IgG (H+L) (115-165-062, Jackson ImmunoResearch), Cy3-conjugated Affinipure goat anti-rat IgG (H+L) (112-165-167, Jackson ImmunoResearch). Sections were washed and mounted with Fluoro-Gel II with DAPI (Cat. 17985-50, Electron microscopy sciences). Sections were imaged by a Zeiss Axio-Imager Z1 with ApoTome attachment or a Leica SP6 inverted confocal microscope as previously described (Salahudeen et al., 2020).

### In vivo organoid transplantation

All animal experimental procedures were approved by an IACUC protocol. Subcutaneous transplantations were performed in NOG-E mice (Taconic) as previously described (Li et al., 2014). Briefly, organoid cells were dissociated into single cell suspensions and mixed in 100 µl of Matrigel and 10^5^-10^6^ cells were subcutaneously injected into the flank. Tumor formation was assessed by palpation and tumors were measured by digital calipers for length, width, and height and ellipsoid volumes were calculated as previously described (Li et al., 2014). Animals were euthanized at predefined endpoints or morbidity criteria and tumor and tissue samples were freshly fixed in formalin and embedded in paraffin.

### In vivo adenoviral injection

10^5^ FGF3 expressing *p53*^*-/-*^ esophageal organoid cells were subcutaneously implanted as above in 20 mice. Mice were monitored for serial tumor measurement and mice with measurable tumors were randomized to achieve a mean of 100 mm^3^ tumor volume. 5×10^8^ pfu of adenovirus expressing FGFR1-ECD-Fc, encoding the soluble extracellular ligand-binding domain of human FGFR1 fused to C-terminal mouse IgG2α Fc (Ad FGFR1-ECD-Fc), or control adenovirus expressing mouse IgG2α Fc (Ad Fc) was injected i.v. retroorbitally and serum was analyzed by Western blot for expression of FGFR1-ECD-Fc or Fc using anti-IgG2α Fc (Jackson ImmunoResearch). All adenoviral inserts were cloned into the E1 region of E1^-^ E3^-^ Ad strain 5 by homologous recombination and amplified in in 293 cells followed by CsCl_2_ gradient purification of virus, as previously described. (Chang et al., 2017; Kuhnert et al., 2010; Kuo et al., 2001; Yan et al., 2017).

### FGF3 ELISA

Organoid conditioned media from log phase growth GFP or FGF3 expressing *p53*^*-/-*^ esophageal organoid cells were assessed according to the manufacturer’s protocol (Aviva Biosciences catalog # OKEH02512).

### In vivo treatment with AZD4547

Mice with subcutaneously either FGF3 or GFP expressing *p53*^*-/-*^ esophageal organoid cells and tumor volumes were serially measured and mice were randomized as above. Mice were treated with either an oral suspension of AZD4547 or vehicle daily as previously described (Gavine et al., 2012) and tumors were serially measured as above.

### Quantitation of CD31+ microvessel density

Immunohistochemistry for CD31 (MAB1398Z, EMD Millipore) was performed on 5 um FFPE tissue sections using Proteinase K antigen retrieval as described (Fairweather et al., 2015). Microvessel density was scored using the Chalkley method as previously described (Fox et al., 1995; Vermeulen et al., 2002). Briefly, FFPE sections of mouse tumors from FGF3 expressing *p53*^*-/-*^ esophageal organoid subcutaneous transplants were subjected to CD31 IHC staining. The mean of three CD31+ hotspots at 20X magnification were scored on the overlap of 25 random points on an ocular grid.

## REFERENCES

Abnet, C.C., Arnold, M., and Wei, W.Q. (2018). Epidemiology of Esophageal Squamous Cell Carcinoma. Gastroenterology 154, 360–373.

Basu, A., and Lambring, C.B. (2021). Akt Isoforms: A Family Affair in Breast Cancer. Cancers (Basel) 13.

Bellacosa, A., de Feo, D., Godwin, A.K., Bell, D.W., Cheng, J.Q., Altomare, D.A., Wan, M., Dubeau, L., Scambia, G., Masciullo, V., et al. (1995). Molecular alterations of the AKT2 oncogene in ovarian and breast carcinomas. Int J Cancer 64, 280–285.

Beroukhim, R., Mermel, C.H., Porter, D., Wei, G., Raychaudhuri, S., Donovan, J., Barretina, J., Boehm, J.S., Dobson, J., Urashima, M., et al. (2010). The landscape of somatic copy-number alteration across human cancers. Nature 463, 899–905.

Boehm, J.S., Garnett, M.J., Adams, D.J., Francies, H.E., Golub, T.R., Hahn, W.C., Iorio, F., McFarland, J.M., Parts, L., and Vazquez, F. (2021). Cancer research needs a better map. Nature 589, 514–516.

Broutier, L., Mastrogiovanni, G., Verstegen, M.M., Francies, H.E., Gavarro, L.M., Bradshaw, C.R., Allen, G.E., Arnes-Benito, R., Sidorova, O., Gaspersz, M.P., et al. (2017). Human primary liver cancer-derived organoid cultures for disease modeling and drug screening. Nat Med 23, 1424–1435.

Cancer Genome Atlas, N. (2015). Comprehensive genomic characterization of head and neck squamous cell carcinomas. Nature 517, 576–582.

Cancer Genome Atlas Research, N., Analysis Working Group: Asan, U., Agency, B.C.C., Brigham Women’s, H., Broad, I., Brown, U., Case Western Reserve, U., Dana-Farber Cancer, I., Duke, U., et al. (2017). Integrated genomic characterization of oesophageal carcinoma. Nature 541, 169–175.

Catenacci, D.V.T., Moya, S., Lomnicki, S., Chase, L.M., Peterson, B.F., Reizine, N., Alpert, L., Setia, N., Xiao, S.Y., Hart, J., et al. (2021). Personalized Antibodies for Gastroesophageal Adenocarcinoma (PANGEA): A Phase II Study Evaluating an Individualized Treatment Strategy for Metastatic Disease. Cancer Discov 11, 308–325.

Chang, J., Mancuso, M.R., Maier, C., Liang, X., Yuki, K., Yang, L., Kwong, J.W., Wang, J., Rao, V., Vallon, M., et al. (2017). Gpr124 is essential for blood-brain barrier integrity in central nervous system disease. Nat Med 23, 450–460.

Curtis, C., Shah, S.P., Chin, S.F., Turashvili, G., Rueda, O.M., Dunning, M.J., Speed, D., Lynch, A.G., Samarajiwa, S., Yuan, Y., et al. (2012). The genomic and transcriptomic architecture of 2,000 breast tumours reveals novel subgroups. Nature 486, 346–352.

Drost, J., van Jaarsveld, R.H., Ponsioen, B., Zimberlin, C., van Boxtel, R., Buijs, A., Sachs, N., Overmeer, R.M., Offerhaus, G.J., Begthel, H., et al. (2015). Sequential cancer mutations in cultured human intestinal stem cells. Nature 521, 43–47.

Du, Y., Li, X., Niu, Q., Mo, X., Qui, M., Ma, T., Kuo, C.J., and Fu, H. (2020). Development of a miniaturized 3D organoid culture platform for ultra-high-throughput screening. J Mol Cell Biol 12, 630–643.

Ebrahimi, M., and Botelho, M. (2017). Adult Stem Cells of Orofacial Origin: Current Knowledge and Limitation and Future Trend in Regenerative Medicine. Tissue Eng Regen Med 14, 719–733.

Fairweather, M., Heit, Y.I., Buie, J., Rosenberg, L.M., Briggs, A., Orgill, D.P., and Bertagnolli, M.M. (2015). Celecoxib inhibits early cutaneous wound healing. J Surg Res 194, 717–724.

Fox, S.B., Leek, R.D., Weekes, M.P., Whitehouse, R.M., Gatter, K.C., and Harris, A.L. (1995). Quantitation and prognostic value of breast cancer angiogenesis: comparison of microvessel density, Chalkley count, and computer image analysis. J Pathol 177, 275–283.

Francies, H.E., Barthorpe, A., McLaren-Douglas, A., Barendt, W.J., and Garnett, M.J. (2019). Drug Sensitivity Assays of Human Cancer Organoid Cultures. Methods Mol Biol 1576, 339–351.

Gavine, P.R., Mooney, L., Kilgour, E., Thomas, A.P., Al-Kadhimi, K., Beck, S., Rooney, C., Coleman, T., Baker, D., Mellor, M.J., et al. (2012). AZD4547: an orally bioavailable, potent, and selective inhibitor of the fibroblast growth factor receptor tyrosine kinase family. Cancer Res 72, 2045–2056.

Gilbert, L.A., Horlbeck, M.A., Adamson, B., Villalta, J.E., Chen, Y., Whitehead, E.H., Guimaraes, C., Panning, B., Ploegh, H.L., Bassik, M.C., et al. (2014). Genome-Scale CRISPR-Mediated Control of Gene Repression and Activation. Cell 159, 647–661.

Guo, X., Wang, X., Wang, Z., Banerjee, S., Yang, J., Huang, L., and Dixon, J.E. (2016). Site-specific proteasome phosphorylation controls cell proliferation and tumorigenesis. Nat Cell Biol 18, 202–212.

H. Pagès, P.A., R. Gentleman, and S. DebRoy. (2021). Biostrings: Efficient manipulation of biological strings. R package version 2.60.2.

Hahn, W.C., Bader, J.S., Braun, T.P., Califano, A., Clemons, P.A., Druker, B.J., Ewald, A.J., Fu, H., Jagu, S., Kemp, C.J., et al. (2021). An expanded universe of cancer targets. Cell 184, 1142–1155.

Hingorani, S.R., Petricoin, E.F., Maitra, A., Rajapakse, V., King, C., Jacobetz, M.A., Ross, S., Conrads, T.P., Veenstra, T.D., Hitt, B.A., et al. (2003). Preinvasive and invasive ductal pancreatic cancer and its early detection in the mouse. Cancer Cell 4, 437–450.

Horlbeck, M.A., Gilbert, L.A., Villalta, J.E., Adamson, B., Pak, R.A., Chen, Y., Fields, A.P., Park, C.Y., Corn, J.E., Kampmann, M., et al. (2016). Compact and highly active next-generation libraries for CRISPR-mediated gene repression and activation. Elife 5.

Huang, L., Holtzinger, A., Jagan, I., BeGora, M., Lohse, I., Ngai, N., Nostro, C., Wang, R., Muthuswamy, L.B., Crawford, H.C., et al. (2015). Ductal pancreatic cancer modeling and drug screening using human pluripotent stem cell- and patient-derived tumor organoids. Nat Med 21, 1364–1371.

Jackson, E.L., Olive, K.P., Tuveson, D.A., Bronson, R., Crowley, D., Brown, M., and Jacks, T. (2005). The differential effects of mutant p53 alleles on advanced murine lung cancer. Cancer Res 65, 10280–10288.

Jones, K.B., Furukawa, S., Marangoni, P., Ma, H., Pinkard, H., D’Urso, R., Zilionis, R., Klein, A.M., and Klein, O.D. (2019). Quantitative Clonal Analysis and Single-Cell Transcriptomics Reveal Division Kinetics, Hierarchy, and Fate of Oral Epithelial Progenitor Cells. Cell Stem Cell 24, 183–192 e188.

Kim, T.M., Yim, S.H., Lee, J.S., Kwon, M.S., Ryu, J.W., Kang, H.M., Fiegler, H., Carter, N.P., and Chung, Y.J. (2005). Genome-wide screening of genomic alterations and their clinicopathologic implications in non-small cell lung cancers. Clin Cancer Res 11, 8235–8242.

Kristensen, V.N., Lingjaerde, O.C., Russnes, H.G., Vollan, H.K., Frigessi, A., and Borresen-Dale, A.L. (2014). Principles and methods of integrative genomic analyses in cancer. Nat Rev Cancer 14, 299–313.

Kryukov, G.V., Wilson, F.H., Ruth, J.R., Paulk, J., Tsherniak, A., Marlow, S.E., Vazquez, F., Weir, B.A., Fitzgerald, M.E., Tanaka, M., et al. (2016). MTAP deletion confers enhanced dependency on the PRMT5 arginine methyltransferase in cancer cells. Science 351, 1214–1218.

Kuhnert, F., Mancuso, M.R., Shamloo, A., Wang, H.T., Choksi, V., Florek, M., Su, H., Fruttiger, M., Young, W.L., Heilshorn, S.C., et al. (2010). Essential regulation of CNS angiogenesis by the orphan G protein-coupled receptor GPR124. Science 330, 985–989.

Kuo, C.J., Farnebo, F., Yu, E.Y., Christofferson, R., Swearingen, R.A., Carter, R., von Recum, H.A., Yuan, J., Kamihara, J., Flynn, E., et al. (2001). Comparative evaluation of the antitumor activity of antiangiogenic proteins delivered by gene transfer. Proc Natl Acad Sci U S A 98, 4605–4610.

Kuuselo, R., Simon, R., Karhu, R., Tennstedt, P., Marx, A.H., Izbicki, J.R., Yekebas, E., Sauter, G., and Kallioniemi, A. (2010). 19q13 amplification is associated with high grade and stage in pancreatic cancer. Genes Chromosomes Cancer 49, 569–575.

Li, X., Francies, H.E., Secrier, M., Perner, J., Miremadi, A., Galeano-Dalmau, N., Barendt, W.J., Letchford, L., Leyden, G.M., Goffin, E.K., et al. (2018). Organoid cultures recapitulate esophageal adenocarcinoma heterogeneity providing a model for clonality studies and precision therapeutics. Nat Commun 9, 2983.

Li, X., Nadauld, L., Ootani, A., Corney, D.C., Pai, R.K., Gevaert, O., Cantrell, M.A., Rack, P.G., Neal, J.T., Chan, C.W., et al. (2014). Oncogenic transformation of diverse gastrointestinal tissues in primary organoid culture. Nat Med 20, 769–777.

Lo, Y.H., Karlsson, K., and Kuo, C.J. (2020). Applications of Organoids for Cancer Biology and Precision Medicine. Nat Cancer 1, 761–773.

Lo, Y.H., Kolahi, K.S., Du, Y., Chang, C.Y., Krokhotin, A., Nair, A., Sobba, W.D., Karlsson, K., Jones, S.J., Longacre, T.A., et al. (2021). A CRISPR/Cas9-engineered ARID1A-deficient human gastric cancer organoid model reveals essential and non-essential modes of oncogenic transformation. Cancer Discov.

Matano, M., Date, S., Shimokawa, M., Takano, A., Fujii, M., Ohta, Y., Watanabe, T., Kanai, T., and Sato, T. (2015). Modeling colorectal cancer using CRISPR-Cas9-mediated engineering of human intestinal organoids. Nat Med 21, 256–262.

Morgan, M., Anders, S., Lawrence, M., Aboyoun, P., Pages, H., and Gentleman, R. (2009). ShortRead: a bioconductor package for input, quality assessment and exploration of high-throughput sequence data. Bioinformatics 25, 2607–2608.

Nadauld, L.D., Garcia, S., Natsoulis, G., Bell, J.M., Miotke, L., Hopmans, E.S., Xu, H., Pai, R.K., Palm, C., Regan, J.F., et al. (2014). Metastatic tumor evolution and organoid modeling implicate TGFBR2 as a cancer driver in diffuse gastric cancer. Genome Biol 15, 428.

Neal, J.T., Li, X., Zhu, J., Giangarra, V., Grzeskowiak, C.L., Ju, J., Liu, I.H., Chiou, S.H., Salahudeen, A.A., Smith, A.R., et al. (2018). Organoid Modeling of the Tumor Immune Microenvironment. Cell 175, 1972–1988 e1916.

Nunez, J.K., Chen, J., Pommier, G.C., Cogan, J.Z., Replogle, J.M., Adriaens, C., Ramadoss, G.N., Shi, Q., Hung, K.L., Samelson, A.J., et al. (2021). Genome-wide programmable transcriptional memory by CRISPR-based epigenome editing. Cell.

Oh, D.Y., and Bang, Y.J. (2020). HER2-targeted therapies - a role beyond breast cancer. Nat Rev Clin Oncol 17, 33–48.

Rueda, O.M., Sammut, S.J., Seoane, J.A., Chin, S.F., Caswell-Jin, J.L., Callari, M., Batra, R., Pereira, B., Bruna, A., Ali, H.R., et al. (2019). Dynamics of breast-cancer relapse reveal late-recurring ER-positive genomic subgroups. Nature 567, 399–404.

Salahudeen, A.A., Choi, S.S., Rustagi, A., Zhu, J., van Unen, V., de la, O.S., Flynn, R.A., Margalef-Català, M., Santos, A.J.M., Ju, J., et al. (2020). Progenitor identification and SARS-CoV-2 infection in human distal lung organoids. Nature 588, 670–675.

Salahudeen, A.A., and Kuo, C.J. (2015). Toward recreating colon cancer in human organoids. Nat Med 21, 215–216.

Sanchez-Garcia, F., Villagrasa, P., Matsui, J., Kotliar, D., Castro, V., Akavia, U.D., Chen, B.J., Saucedo-Cuevas, L., Rodriguez Barrueco, R., Llobet-Navas, D., et al. (2014). Integration of genomic data enables selective discovery of breast cancer drivers. Cell 159, 1461–1475.

Sato, T., Stange, D.E., Ferrante, M., Vries, R.G., Van Es, J.H., Van den Brink, S., Van Houdt, W.J., Pronk, A., Van Gorp, J., Siersema, P.D., et al. (2011). Long-term expansion of epithelial organoids from human colon, adenoma, adenocarcinoma, and Barrett’s epithelium. Gastroenterology 141, 1762–1772.

Taira, N., Nihira, K., Yamaguchi, T., Miki, Y., and Yoshida, K. (2007). DYRK2 is targeted to the nucleus and controls p53 via Ser46 phosphorylation in the apoptotic response to DNA damage. Mol Cell 25, 725–738.

Tandon, V., de la Vega, L., and Banerjee, S. (2021). Emerging roles of DYRK2 in cancer. J Biol Chem 296, 100233.

Tomlins, S.A., Rhodes, D.R., Perner, S., Dhanasekaran, S.M., Mehra, R., Sun, X.W., Varambally, S., Cao, X., Tchinda, J., Kuefer, R., et al. (2005). Recurrent fusion of TMPRSS2 and ETS transcription factor genes in prostate cancer. Science 310, 644–648.

Turner, N., Pearson, A., Sharpe, R., Lambros, M., Geyer, F., Lopez-Garcia, M.A., Natrajan, R., Marchio, C., Iorns, E., Mackay, A., et al. (2010). FGFR1 amplification drives endocrine therapy resistance and is a therapeutic target in breast cancer. Cancer Res 70, 2085–2094.

Valenzuela, C.A., Vargas, L., Martinez, V., Bravo, S., and Brown, N.E. (2017). Palbociclib-induced autophagy and senescence in gastric cancer cells. Exp Cell Res 360, 390–396.

van de Wetering, M., Francies, H.E., Francis, J.M., Bounova, G., Iorio, F., Pronk, A., van Houdt, W., van Gorp, J., Taylor-Weiner, A., Kester, L., et al. (2015). Prospective derivation of a living organoid biobank of colorectal cancer patients. Cell 161, 933–945.

van Dyk, E., Hoogstraat, M., Ten Hoeve, J., Reinders, M.J., and Wessels, L.F. (2016). RUBIC identifies driver genes by detecting recurrent DNA copy number breaks. Nat Commun 7, 12159.

Vermeulen, P.B., Gasparini, G., Fox, S.B., Colpaert, C., Marson, L.P., Gion, M., Belien, J.A., de Waal, R.M., Van Marck, E., Magnani, E., et al. (2002). Second international consensus on the methodology and criteria of evaluation of angiogenesis quantification in solid human tumours. Eur J Cancer 38, 1564–1579.

Weaver, A., and Bossaer, J.B. (2021). Fibroblast growth factor receptor (FGFR) inhibitors: A review of a novel therapeutic class. J Oncol Pharm Pract 27, 702–710.

Wong, G.S., Zhou, J., Liu, J.B., Wu, Z., Xu, X., Li, T., Xu, D., Schumacher, S.E., Puschhof, J., McFarland, J., et al. (2018). Targeting wild-type KRAS-amplified gastroesophageal cancer through combined MEK and SHP2 inhibition. Nat Med 24, 968–977.

Yan, K.S., Janda, C.Y., Chang, J., Zheng, G.X.Y., Larkin, K.A., Luca, V.C., Chia, L.A., Mah, A.T., Han, A., Terry, J.M., et al. (2017). Non-equivalence of Wnt and R-spondin ligands during Lgr5(+) intestinal stem-cell self-renewal. Nature 545, 238–242.

Yang, X., Boehm, J.S., Yang, X., Salehi-Ashtiani, K., Hao, T., Shen, Y., Lubonja, R., Thomas, S.R., Alkan, O., Bhimdi, T., et al. (2011). A public genome-scale lentiviral expression library of human ORFs. Nat Methods 8, 659–661.

Zack, T.I., Schumacher, S.E., Carter, S.L., Cherniack, A.D., Saksena, G., Tabak, B., Lawrence, M.S., Zhsng, C.Z., Wala, J., Mermel, C.H., et al. (2013). Pan-cancer patterns of somatic copy number alteration. Nat Genet 45, 1134–1140.

