## Supplementary material for "Pan-cancer organoid validation of tumor outlier chromosomal amplification events": Key Resources Table.pdf

| REAGENT or RESOURCE | SOURCE | IDENTIFIER |
|---|---|---|
| **Antibodies** | | |
| DYRK2 antibody | Cell Signaling | 8143 |
| GAPDH (D16H11) Rabbit mAb | Cell Signaling | 5174S |
| Peroxidase AffiniPure Goat Anti-Rabbit IgG (H+L) | Jackson ImmunoResearch | 711-035-152 |
| Mouse anti-E-cadherin | BD Biosciences | BD610182 |
| Alexa fluor 647 labeled-anti-CK5 | Abcam | ab193895 |
| Rat-anti-Ki67 | eBiosciences | 14-5698-82 |
| Cy3-conjugated affinipure goat anti-mouse IgG (H+L) | Jackson ImmunoResearch | 115-165-062 |
| Cy3-conjugated affinipure goat anti-rat IgG (H+L) | Jackson ImmunoResearch | 112-165-167 |
| Peroxidase AffiniPure Goat Anti-Mouse IgG, Fcγ subclass 2a specific | Jackson ImmunoResearch | 115-035-206 |
| **Bacterial and virus strains** | | |
| Lentivirus for p53 R175H | Ken Scott laboratory | KS5 |
| Ad Cre GFP | University of Iowa Vector Core | 1174 |
| Adenovirus Fc | Stanford University | N/A |
| DYRK2 lentivirus | Genetic Perturbation Platform - Broad Institute | TRCN489007 |
| CCSB-Broad Lentiviral Expression Library | The Broad ORF collection | N/A |
| **Biological samples** | | |
| $p53^{R175H}$ gastric organoids | Stanford University | N/A |
| $Kras^{G12D}$ pancreatic organoids | Stanford University | N/A |
| Human $APC^{-/-}$ Colon Organoids | Stanford University | N/A |
| $p53^{-/-}$ oral squamous cell carcinoma organoids | Stanford University | N/A |
| $p53^{-/-}$ esophageal squamous cell carcinoma organoids | Stanford University | N/A |
| $Kras^{G12D}$ $p53^{-/-}$ lung adenocarcinoma | Stanford University | N/A |
| oral mucosa (OM) mouse organoids | Stanford University | N/A |
| **Chemicals, peptides, and recombinant proteins** | | |
| cOmplete™, Mini Protease Inhibitor Cocktail | Sigma Aldrich | 4693124001 |
| SuperSignal™ West Pico PLUS Chemiluminescent Substrate | ThermoFisher Scientific | 34580 |
| Fluoro-Gel II with DAPI | Electron Microscopy Sciences | 17985-50 |
| 1mM HEPES | Thermo Fisher Scientific | BP299100 |
| Nicotinomide | Sigma | N0636-500G |
| GlutaMAX™ Supplement | ThermoFisher Scientific | 31980030 |
| 1mM N-acetyl-l-cysteine | Sigma-Aldrich | A7250-100G |

| | | |
|---|---|---|
| B27 | Invitrogen | 17504-001 |
| A83-01 | Tocris | 2939 |
| Recombinant human EGF | Peprotech | AF-100-15 |
| Recombinant human Noggin | Peprotech | 120-10-C |
| 2-mercaptoethanol | ThermoFisher Scientific | 21985023 |
| DTT | Sigma | 10197777001 |
| Penicillin-Streptomycin-Glutamine (100X) | Life Technologies | 10378016 |
| **Critical commercial assays** | | |
| RNeasy Plus Mini Kit | Qiiagen | 74104 |
| iScript Reverse Transcription Supermix | BioRad | 1708841 |
| FGF3 ELISA | Aviva Systems Biology | OKEH02512 |
| Hs FGF3 Taqman Assay | ThermoFisher | 4448892 |
| Hs beta actin Taqman Assay | ThermoFisher | 401846 |
| **Deposited data** | | |
| Outlier Barcode Sequencing Data | CTDD Portal | Pending |
| TCGA Expression Data | Firehose, Broad Institute | http://gdac.broadinstitute.org/ |
| Broad Genomic Perturbation Platform | Broad Institute | https://portals.broadinstitute.org/gpp/public/ |
| **Experimental models: Cell lines** | | |
| HEK293T | ATCC | CRL-11268 |
| **Experimental models: Organisms/strains** | | |
| Il2rgtm1Sug/JicTac (NOG) Immunodeficient mice | Taconic | NOG-F |
| C57BL/6NTac mice | Taconic | B6-F |
| p53 flox/flox mice | Jackson Labs | 008462 |
| LSL-Kras G12D mice | Jackson Labs | 008179 |
| **Oligonucleotides** | | |
| *Tp53* Forward Primer: tgaggttcgtgtttgtgcct | Stanford PAN Facility | N/A |
| *Tp53* Reverse Primer: gcagttcagggcaaaggact | Stanford PAN Facility | N/A |
| *Ki67* Forward Primer: agctctgcagtctccacaac | Stanford PAN Facility | N/A |
| *Ki67* Reverse Primer: ctctgcctcgtgactgtgtt | Stanford PAN Facility | N/A |
| *Cdkn1a* Forward Primer: ttgtcgctgtcttgcactct | Stanford PAN Facility | N/A |
| *Cdkn1a* Reverse Primer: tttcggccctgagatgttcc | Stanford PAN Facility | N/A |
| *DYRK2* Forward Primer: cagtgctcacgacacaacca | Stanford PAN Facility | N/A |
| *DYRK2* Reverse Primer: ccgtctatgaatgctgtccag | Stanford PAN Facility | N/A |
| *Gapdh* Forward Primer: tgaacgggaagctcactgg | Stanford PAN Facility | N/A |
| *Gapdh* Reverse Primer: tccaccaccctgttgctgta | Stanford PAN Facility | N/A |
| APC gRNA: | | |
| **Recombinant DNA** | | |
| px330 CRISPR Plasmid | AddGene | 42230 |
| **Software and algorithms** | | |

| | | |
|---|---|---|
| Excel | Microsoft | https://www.microsoft.com/en-us/microsoft-365/excel |
| Powerpoint | Microsoft | https://www.microsoft.com/en-us/microsoft-365/powerpoint |

| | | |
|---|---|---|
| Prism 8.2.0 | GraphPad | https://www.graphpad.com |
| RStudio | RStudio | https://rstudio.com |
| **Other** | | |
| LUCENTBLUE X-RAY FILM, 8x10cm FOR CHEMILUMINESCENCE | E&K Scientific | EK-5129 |
| iScript™ Reverse Transcription Supermix | Bio-Rad | 1708841 |
| Power SYBR™ Green PCR Master Mix | Thermofisher Scientific | 4367659 |
| Cultrex Rat Collagen I, Lower Viscosity (3 mg/ml) | Trevigen | 3443-100-01 |
| AlamarBlue Cell Viability Reagent | Life Technologies | DAL1100 |
| Citrate Buffer, pH 6.0, Antigen Retrieval Solution | Millipore Sigma | C9999 |
| Normal donkey serum | Jackson ImmunoResearch | 017-000-121 |
| Trevigen Cultrex Reduced Growth Factor BME Type 2 | R&D Systems | 3533-005-02 |
| RIPA Buffer | Thermofisher Scientific | 89900 |
| 4%–12% Bis-Tris Gels | Thermofisher Scientific | |
| TrypLE™ Express (1X), no Phenol Red | Life Technologies | 12604021 |
| DNase I | Worthington | LS006328 |
| Collagenase Type 4 5x50mg | Worthington | LS004212 |
| DNeasy Blood and Tissue Kit | Qiagen | 69506 |
| Polybrene Infection / Transfection Reagent | Sigma | TR-1003-G |
