## Supplementary material for "Pan-cancer organoid validation of tumor outlier chromosomal amplification events": Salahudeen_etal_Supplemental_Figs.pdf

### Figure S1

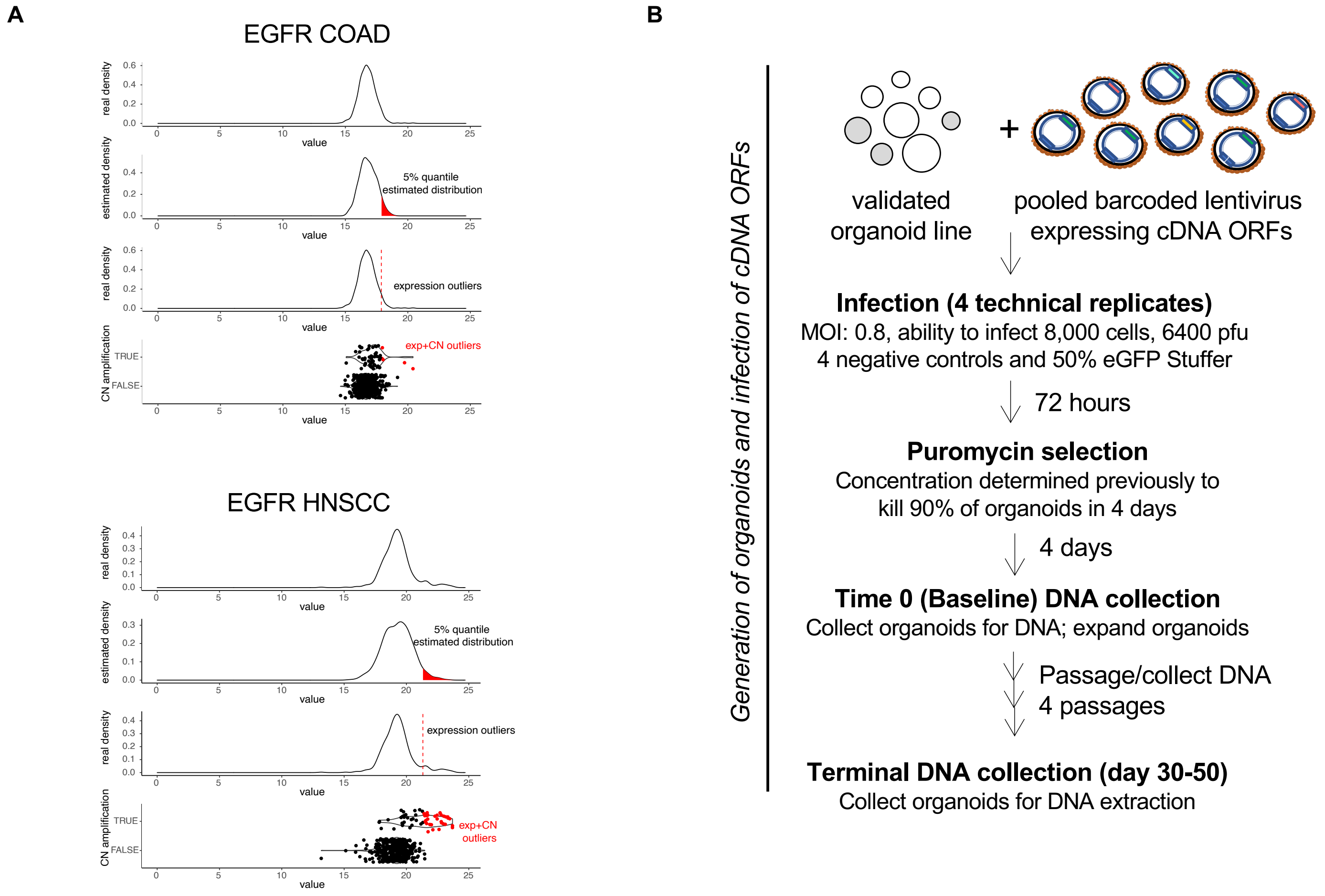


**Figure S1. Identification and Validation of candidate SCNA drivers (related to Fig. 1).**

A) Schematic of integrative analysis to nominate outlier gene candidates for EGFR in colon adenocarcinoma (COAD) (top) and head and neck squamous carcinoma (HNSCC) (bottom). Top panel shows the expression counts density for EGFR to obtain the mean and standard deviation from these distributions. In the second panel, using the mean and standard deviation, theoretical normal distributions were generated and the 5% quantile of higher expression (red colored distribution) was used to define a threshold that was later applied to the original distribution (red line, third panel). Samples with expression higher than these thresholds were classified as expression outliers. In the last panel the boxplots shows the distribution of expression across copy number amplified and not amplified samples. The final outlier list is the intersection between the amplified samples and the expression outliers and the amplification in the COAD samples lack an effect in the expression, thus most of the amplified samples are not classified as outliers. In contrast, in HNSC (bottom panels) there is a significant number of CN amplified samples with a subsequent change in expression resulting in greater number of amplified samples being classified as outliers.

B) Schematic of pooled ORF screening experiments in this study.

### Figure S2

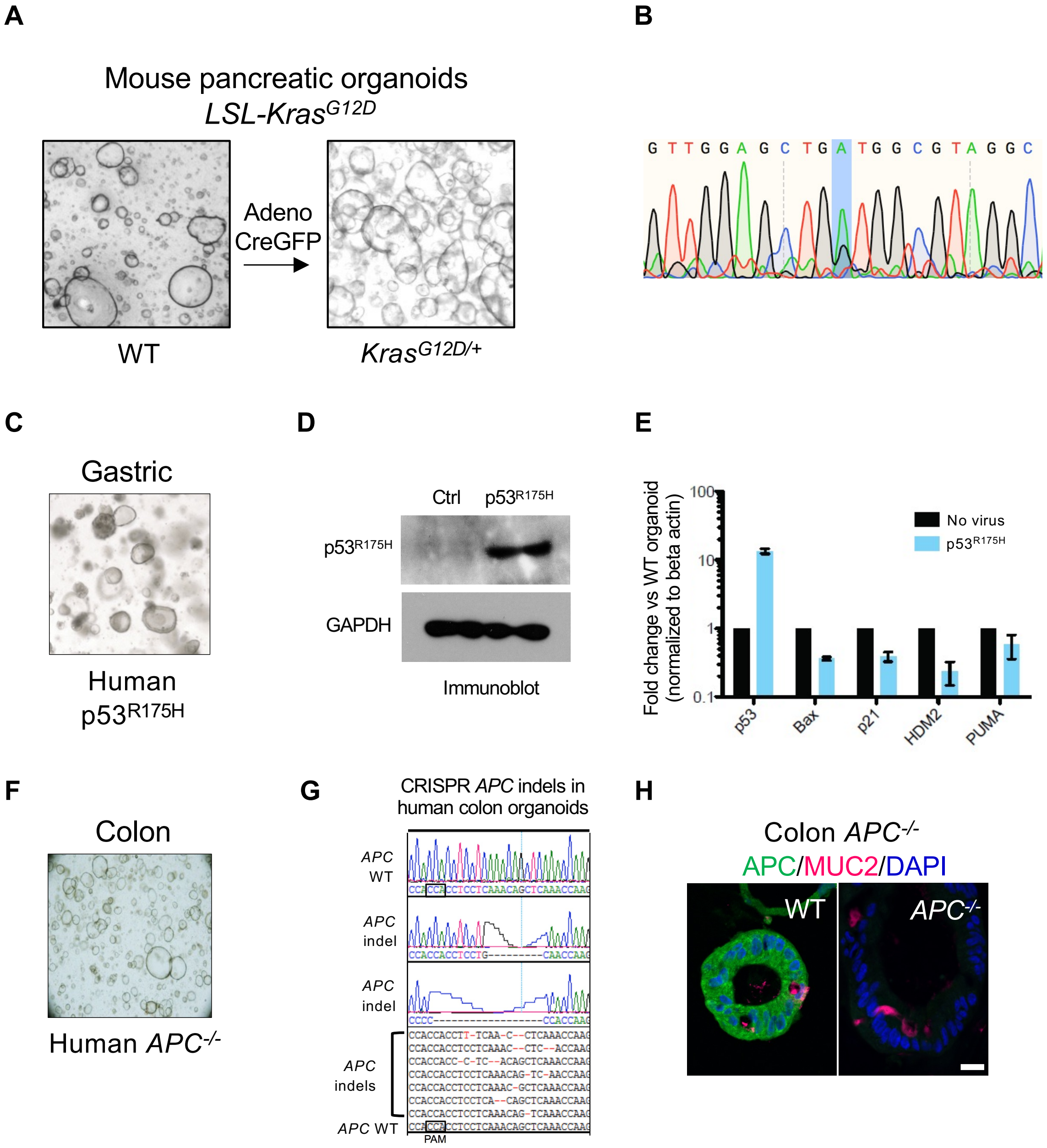


**Figure S2. Generation and validation of tissue contextual models of pancreatic, gastric, and colon carcinomas (related to Fig. 1)**

A) Schematic of *Kras$^{G12D}$* pancreatic organoid derivation.
B) Sanger sequencing of *Kras* cDNA.
C) Brightfield microscopy of human gastric *p53$^{R175H}$* organoids.
D) Immunoblotting of p53$^{R175H}$ in transduced versus control gastric organoids.
E) qRT-PCR of *p53* target genes in transduced versus control gastric organoids.
F) Brightfield microscopy of human colon *APC$^{-/-}$* organoids.
G) Sanger tracing of TA cloned genomic PCR amplicons of the CRISPR KO *APC* locus demonstrating indel mutagenesis.
H) Immunofluorescence of wild type versus *APC* CRISPR KO human colon organoids.

### Figure S3

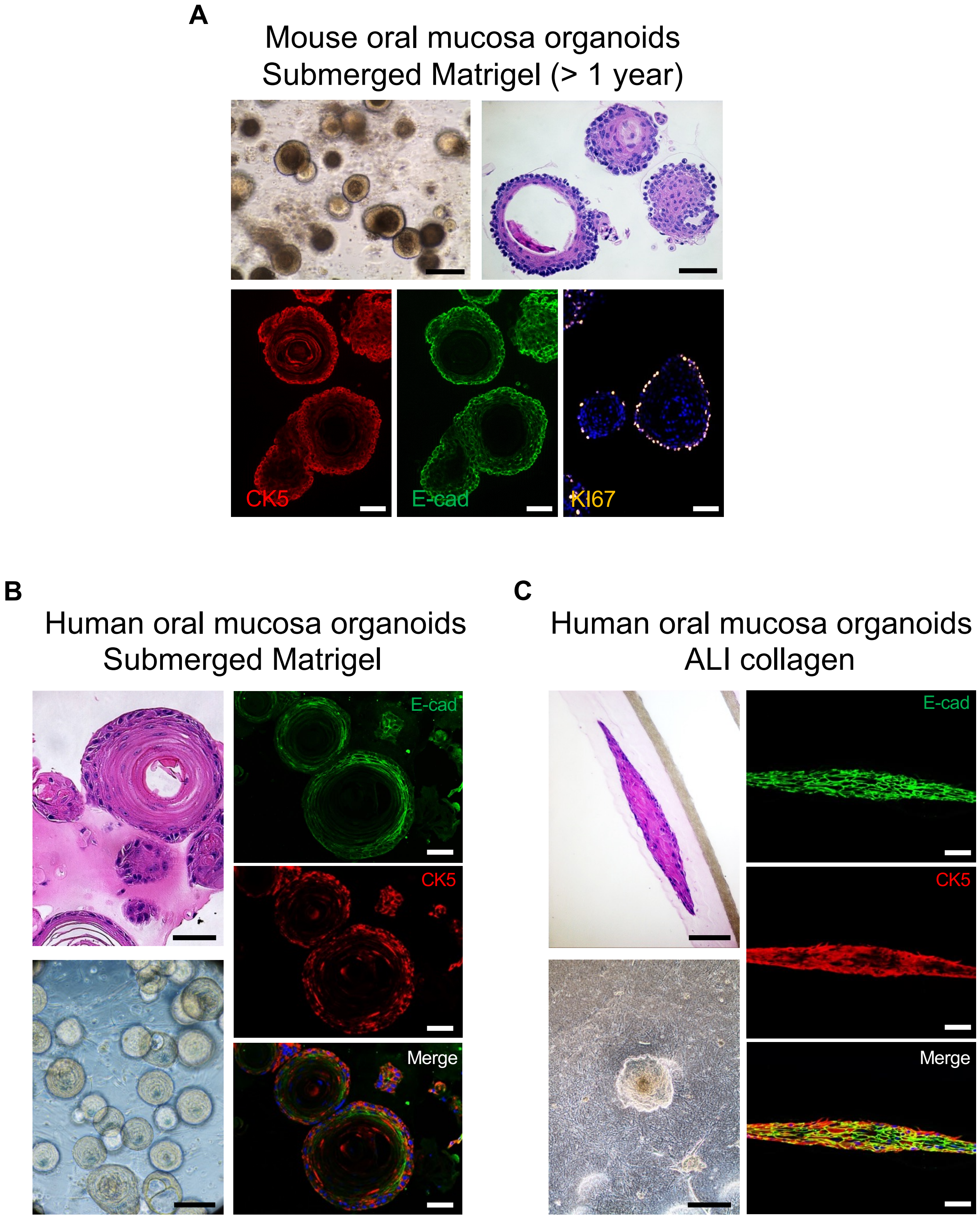


**Figure S3. Evaluation of murine and oral mucosal organoids in submerged Matrigel and ALI (related to Fig. 2)**
A) Submerged Matrigel cultures of wild type murine oral mucosa organoids at day 9, scale bars: brightfield; 200 µm, H&E staining; 50 µm, IF staining; 50 µm.
B) Submerged Matrigel cultures of normal human oral mucosa at day 13, scale bars: H&E staining; 50 µm, brightfield; 100 µm, IF staining; 50 µm.
C) Air-liquid interface cultures of normal human oral mucosa at day 18, scale bars: H&E staining; 100 µm, brightfield; 500 µm, IF staining; 50 µm.

### Figure S4

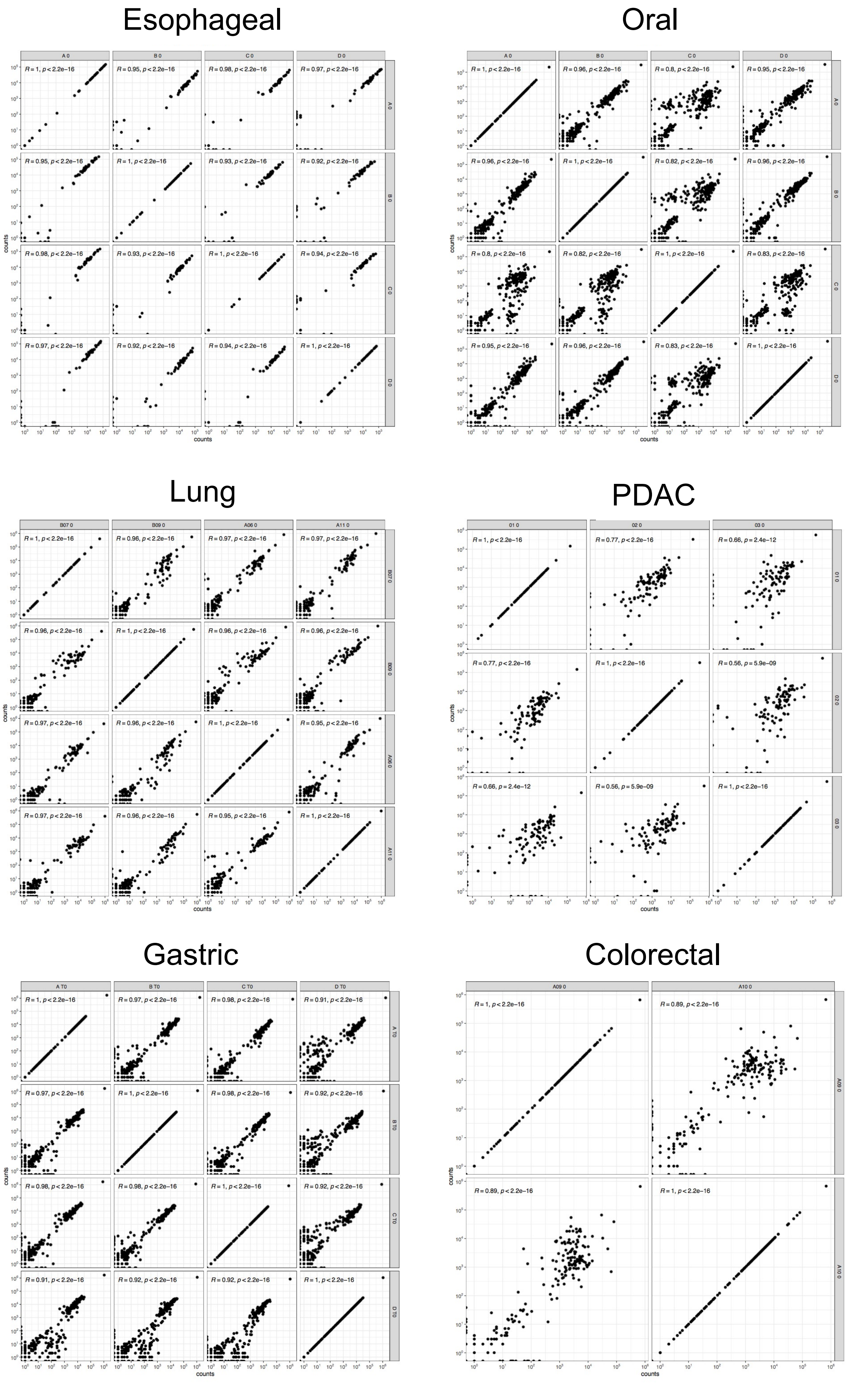


**Figure S4. Scatter plots correlating Next Generation Sequencing barcode counts of time point zero technical replicates (related to Fig. 5)**

### Figure S5

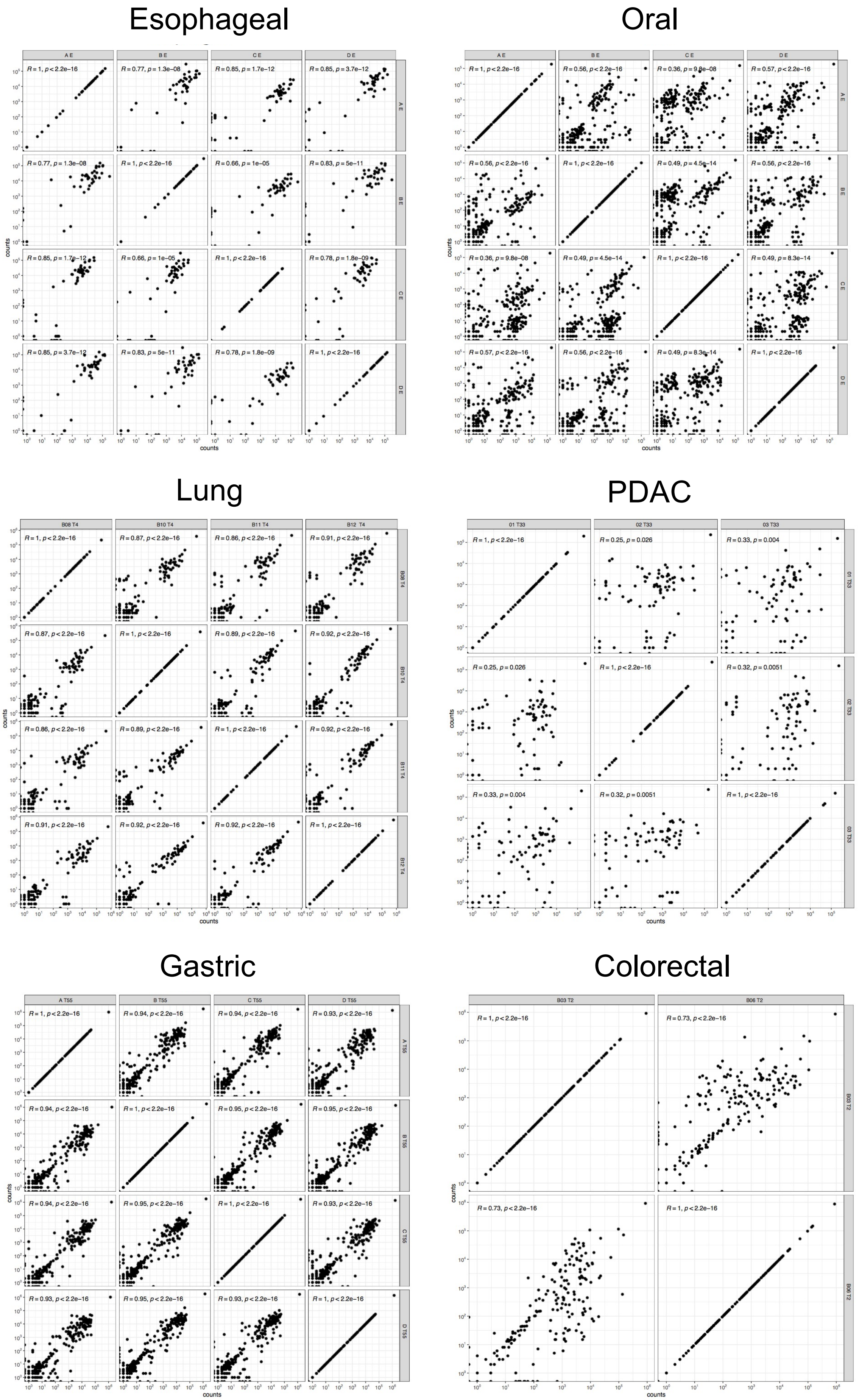


**Figure S5. Scatter plots correlating Next Generation Sequencing barcode counts of terminal time point technical replicates (related to Fig. 5)**

### Figure S6

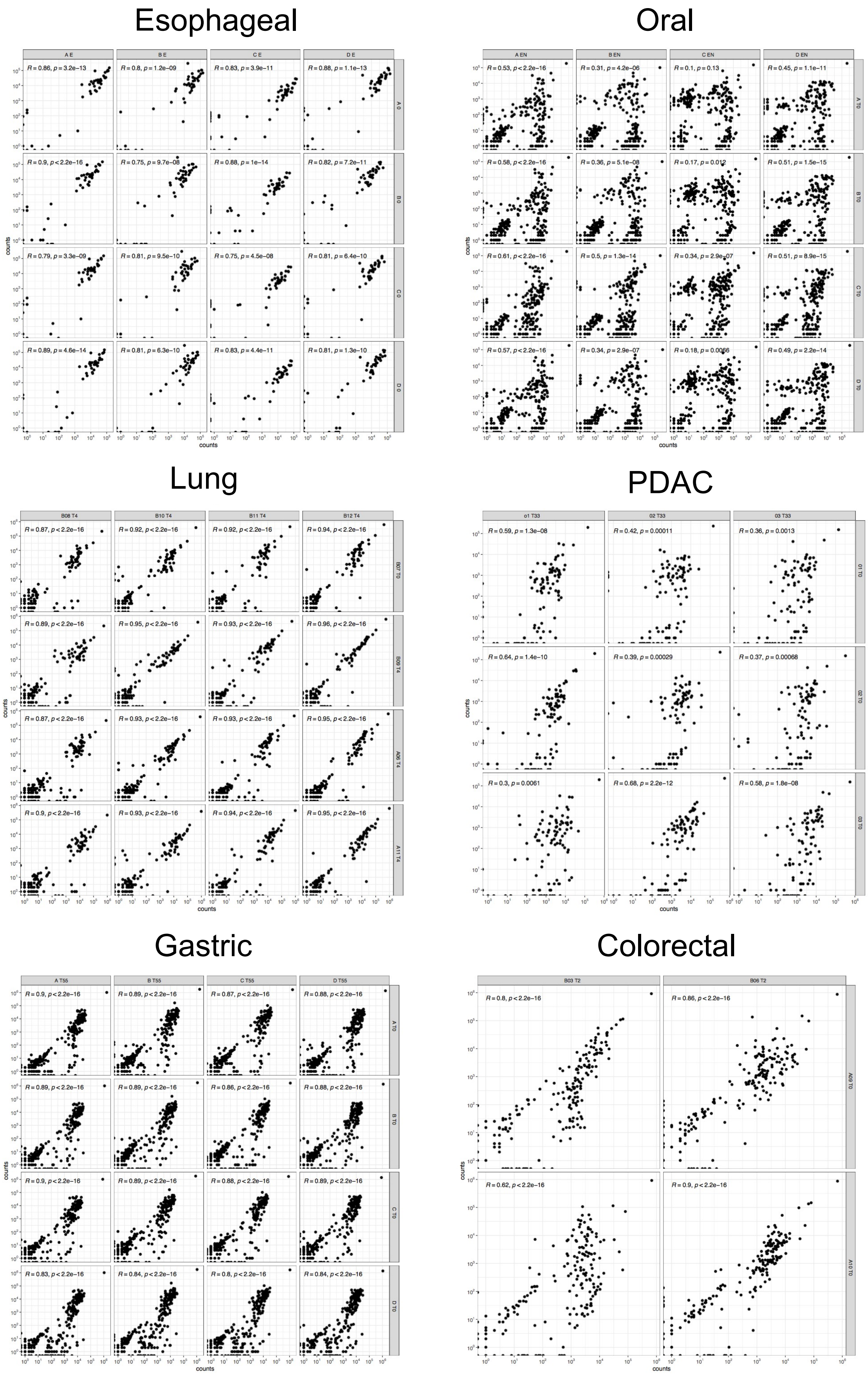


**Figure S6. Scatter plots correlating Next Generation Sequencing barcode counts of terminal time points versus time point zero technical replicates (related to Fig. 5).**

### Figure S7

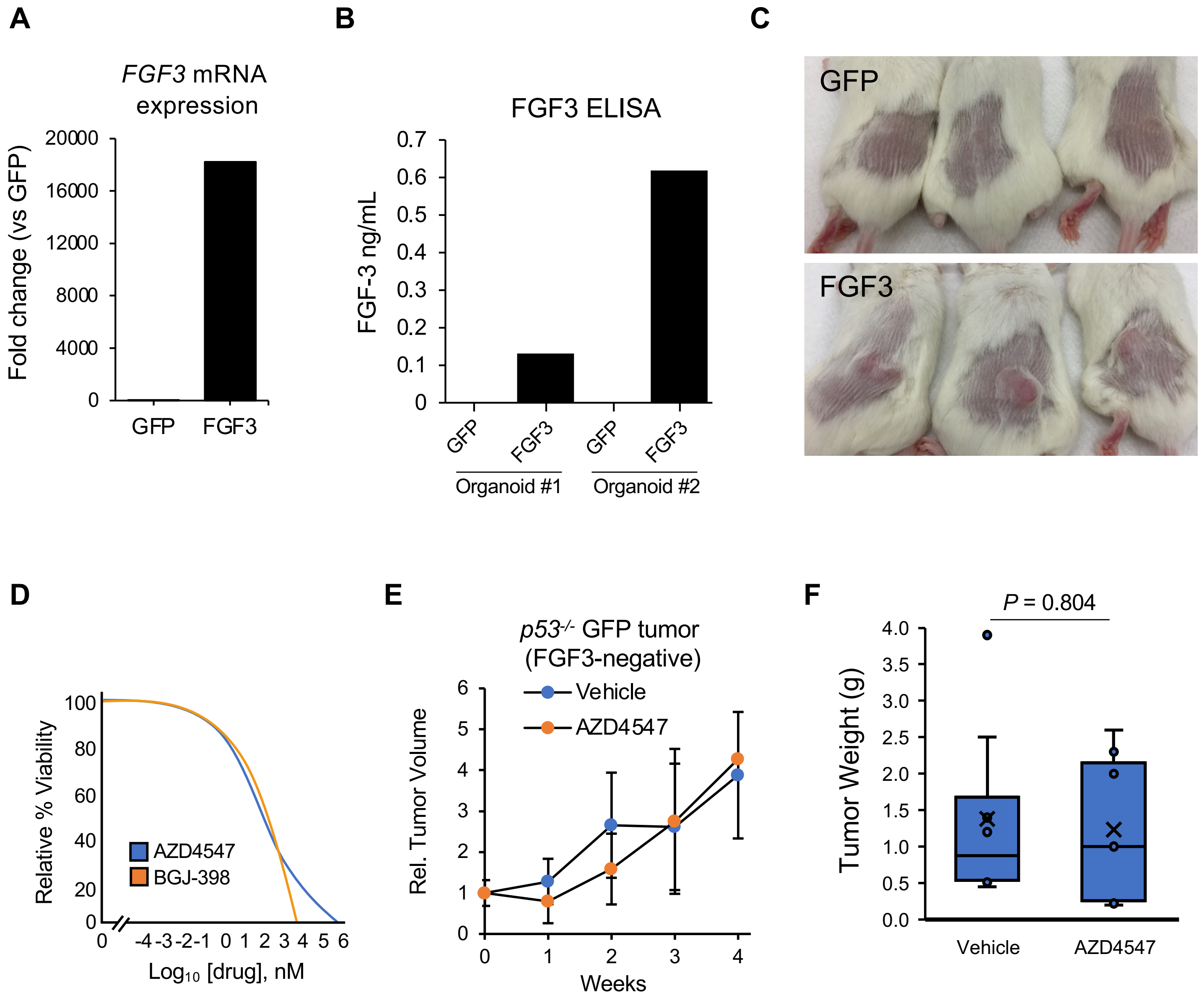


**Figure S7. Evaluation of mechanisms of FGF3 mediated tumorigenesis in *p53*$^{-/-}$ esophageal organoids (related to Fig. 7)**

A) qRT-PCR of FGF3 mRNA in GFP and FGF3 expressing *p53*$^{-/-}$ esophageal organoids.

B) Bar plots of FGF3 ELISA measurements in organoid conditioned media.

C) Photographs of subcutaneous tumor formation of GFP expressing and FGF3 expressing *p53*$^{-/-}$ esophageal organoids.

D) Fitted dose response curves of BGJ398 ($EC_{50}$ = 567 nM) and AZD4547 ($EC_{50}$ = 348 nM) FGFR inhibitors on FGF3-overexpressing *p53*$^{-/-}$ esophageal organoids. Each group had n = 3 technical replicates.

E) Serial tumor measurements of GFP expressing *p53*$^{-/-}$ esophageal organoids treated with vehicle or AZD4547. Each group had n = 10 biological replicates.

F) Terminal tumor weights of D.
